# β-aminopropionitrile Induces Distinct Pathologies in the Ascending and Descending Thoracic Aortic Regions of Mice

**DOI:** 10.1101/2023.10.22.563474

**Authors:** Michael K. Franklin, Hisashi Sawada, Sohei Ito, Deborah A. Howatt, Naofumi Amioka, Ching-Ling Liang, Nancy Zhang, David B. Graf, Jessica J. Moorleghen, Yuriko Katsumata, Hong S. Lu, Alan Daugherty

## Abstract

**BACKGROUND:** β-aminopropionitrile (BAPN) is a pharmacological inhibitor of lysyl oxidase and lysyl oxidase-like proteins. Administration of BAPN promotes aortopathies, although there is a paucity of data on experimental conditions to generate pathology. The objective of this study was to define experimental parameters and determine whether equivalent or variable aortopathies were generated throughout the aortic tree during BAPN administration in mice.

**METHODS:** BAPN was administered in drinking water for a period ranging from 1 to 12 weeks. The impacts of BAPN were first assessed with regard to dose, strain, age, and sex. BAPN-induced aortic pathological characterization was conducted using histology and immunostaining. To investigate the mechanistic basis of regional heterogeneity, ascending and descending thoracic aortas were harvested after one week of BAPN administration before the appearance of overt pathology.

**RESULTS:** BAPN-induced aortic rupture predominantly occurred or originated in the descending thoracic aorta in young C57BL/6J or N mice. No apparent differences were found between male and female mice. For mice surviving 12 weeks of BAPN administration, profound dilatation was consistently observed in the ascending region, while there were more heterogeneous changes in the descending thoracic region. Pathological features were distinct between the ascending and descending thoracic regions. Aortic pathology in the ascending region was characterized by luminal dilatation and elastic fiber disruption throughout the media. The descending thoracic region frequently had dissections with false lumen formation, collagen deposition, and remodeling of the wall surrounding the false lumen. Cells surrounding the false lumen were predominantly positive for α-smooth muscle actin. One week of BAPN administration compromised contractile properties in both regions equivalently, and RNA sequencing did not show obvious differences between the two aortic regions in smooth muscle cell markers, cell proliferation markers, and extracellular components.

**CONCLUSIONS:** BAPN-induced pathologies show distinct, heterogeneous features within and between ascending and descending aortic regions in mice.

## INTRODUCTION

Aortopathies represent a spectrum of pathologies including luminal dilations, medial dissections, transmural rupture, and pathological remodeling of the aortic wall.^1^ Given the increased focus on the devastating outcomes of aortopathies, there has been an enhanced interest in understanding the pathological features of this disease. A number of animal models have been developed using genetic and chemical manipulations to study the pathogenesis and molecular mechanisms of aortopathies. One mouse model used for studying aortopathies is β-aminopropionitrile (BAPN) administration into 3-4-week-old C57BL/6J mice.

Loss of the integrity of elastic fibers is a common feature of aortopathies.^2, 3^ Formation and maturation of elastic fibers is a complex and undefined process in which the stabilization of elastic fibers relies on the crosslinking of four vicinal lysine residues to form desmosine in a reaction catalyzed by lysyl oxidase (LOX) and LOX-like proteins.^4, 5^ BAPN is a pharmacological inhibitor of LOX and LOX-like proteins. Administration of BAPN has long been recognized for promoting aortopathies.^6^ Aortic rupture during BAPN administration was noted in young rats and turkeys in the 1950s.^7, 8^ BAPN was subsequently used in mice in conjugation with angiotensin II infusion subcutaneously to produce aortopathies in adult mice,^9^ which has been replicated by multiple groups.^10, 11^ Further studies demonstrated that BAPN administration alone promotes aortopathies in young C57BL/6 mice before the maturation of elastic fibers.^12^

Although BAPN-induced aortic pathologies are predominant in the thoracic region,^6^ pathological features in this aortic region have not been extensively investigated. Most studies reported overt pathological changes in 3-4-week-old male C57BL/6 mice administered BAPN for 4 weeks.^6, 12–16^ Only two mouse strains, C57BL/6 and FVB, have been compared for the susceptibility of BAPN-induced aortopathies.^12^ The reports on BAPN-induced aortic pathologies have not distinguished between the J and N substrains of C57BL/6, although there is evidence of differences between the two substrains in angiotensin II-induced aortopathy.^17^ In the present study, we performed a dose-response curve in young C57BL/6J mice and compared BAPN-induced aortopathies in five mouse strains that have been used commonly for genetic manipulations. We also compared the incidence of aortopathies in both male and female young versus mature C57BL/6J mice. We characterized pathological features in both the ascending and descending thoracic aortic regions at their acute phases (1-4 weeks of BAPN administration) and an advanced stage (12 weeks of BAPN administration) in young male C57BL/6J mice. We noted that BAPN-induced aortic pathologies were highly heterogeneous. While the aortopathies were restricted to the thoracic regions, distinct characteristics of aortic pathologies between the ascending and descending thoracic regions were evident.

## MATERIALS AND METHODS

### Mice

Mice of the following strains were purchased from The Jackson Laboratory: C57BL/6J (# 000664), C57BL/6N (# 005304), B6/129SF1 (# 101043), 129X1 (# 000691), and FVB (# 001800). Study mice were maintained in individually vented cages (5 mice/cage) on a light : dark cycle of 14 : 10 hours. Teklad Sani-Chip (Cat # 7090A; Inotiv) was used as cage bedding. Mice were fed a normal rodent laboratory diet (Diet # 2918; Inotiv) and given drinking water alone (control) or drinking water containing BAPN ad libitum. Detailed mouse strain information and inclusion and exclusion criteria are provided in Supplemental Materials Major Resources Tables. All mouse experiments were conducted in accordance with the ARRIVE guidelines (Animal Research: Reporting of In Vivo Experiments) and were performed with the approval of the University of Kentucky Institutional Animal Care and Use Committee.

### BAPN Administration

BAPN (0.1%, 0.3%, or 0.5% wt/vol; CAS: 2079-89-2; Cat # A3134-25G, Millipore-Sigma or Cat # A0796-500G, TCI Co.) was administered via drinking water when mice were 3-4 or 26 weeks of age. Fresh drinking water, with or without BAPN, was replaced twice each week.

### Necropsy

All study mice were checked at least once every day. Necropsies were performed immediately to determine the cause of death after carcasses were found. Aortic rupture was defined as the presence of extravascular blood that accumulated in a body cavity. The location of blood egress was determined by the location of the blood clot and a discernable disruption of the aortic wall.

### Ultrasound measurements

Ultrasonography was performed by one investigator, who was blinded to the mouse identity, using our standardized protocols.^18^ Briefly, mice were anesthetized using inhaled isoflurane (2-3% vol/vol) and maintained at a heart rate of >400 beats per minute during image capture to reduce anesthesia exposure and maintain consistent heart rate between animals (SomnoFlo, Kent Scientific). The order by which mice were subject to ultrasound was randomized. Ultrasound images were captured in the right parasternal view using a Vevo 3100 system with a 40 MHz transducer (VisualSonics, Fujifilm). Images captured were standardized according to two anatomical landmarks: the innominate artery branch point and aortic valves. Aortic lumen was defined as edge-to-edge of the ascending aortic wall between the sinotubular junction and the innominate artery. The largest luminal diameters of ascending aortas were measured in end-diastole over three cardiac cycles. The measurements were performed by an investigator who did not capture the images and was blinded to study groups. Some measurements were performed by two investigators independently to compare the consistency of measurements.

### Quantification of aortic diameters using in situ images

Mice surviving 12 weeks of BAPN administration were euthanized by a cocktail of ketamine (90 mg/kg) and xylazine (10 mg/kg). The right atrial appendage was excised and saline (∼10 ml) was perfused via the left ventricle. Periaortic tissues were carefully dissected out and a black plastic sheet was inserted underneath the aorta to improve image contrast.^19, 20^ A millimeter ruler was placed next to the aorta for calibration of measurements. In situ aortic images were captured with a Nikon SMZ (25 or 800) stereoscope (Nikon). Aortic images were analyzed using NIS-Elements AR software (Version 5.11, Nikon). The measurement software was calibrated using the ruler on each image. To measure aortic diameters, a measurement line was drawn perpendicularly to the aortic axis at the most dilated area of each thoracic aortic region (ascending/arch and descending thoracic regions). Measurements were verified by an individual who was blinded to the study groups.

### Histology and immunostaining

For histological analyses, ascending and descending thoracic aortas were harvested from surviving mice after 2, 4, or 12 weeks of either vehicle or BAPN administration and immersed in buffered formalin (10% wt/vol), followed by incubation in ethanol (70% vol/vol), and subsequently embedded in paraffin. Paraffin-embedded sections (5 µm) were deparaffinized using limonene (Cat #183164, Millipore-Sigma). Hematoxylin and eosin (H&E, Cat # 26043-06, Electron Microscopy Sciences, Cat # AB246824, abcam), Verhoeff iron hematoxylin, and Movat’s pentachrome staining (Cat # k042, Poly Scientific R&D) were performed. Immunostaining was performed using primary and secondary antibodies listed in Supplemental Materials Major Resources Tables. NovaRed (Cat #SK-4805, Vector) or AEC (Cat #SK-4205, Vector) were used as chromogens. Images of histological staining and immunostaining were captured using an Axioscan Z1 or 7 (Zeiss) and imaged using ZEN v3.1 blue edition (Zeiss).

### Isometric force analysis of ascending and descending thoracic aortas

Mice were euthanized by ketamine:xylazine anesthesia and aortas were placed in oxygenated (5% CO_2_ and 95% O_2_) modified Krebs–Henseleit physiological salt solution (PSS, in mM: NaCl 130, NaHCO_3_ 24.9, KCl 4.7, KH_2_PO_4_ 1.18, CaCl_2_ 1.6, MgSO_4_ 1.17, glucose 5.5, and EDTA 0.026). Sections (2 mm) of the ascending and proximal descending aorta were mounted on pins (200 µm) in DMT myograph chambers (DMT630MA). Vessels were equilibrated at normal physiological conditions and underwent a normalization procedure to determine the optimal passive tension of each vessel segment. The viability of the aortic segments was determined using high potassium physiological salt solution (KPSS, inLmM: NaCl 74.7, NaHCO_3_ 24.9, KCl 60, KH_2_PO_4_ 1.18, CaCl_2_ 1.6, MgSO_4_ 1.17, glucose 5.5, and EDTA 0.026). Concentration-response curves were obtained in tissues initially contracted with 5-hydroxytryptamine (-9 to -6 logM). The muscle tension generated was plotted based on the force (mN) generated with each concentration.

### RNA sequencing of the ascending and descending thoracic aortas

The ascending aorta and the descending thoracic aorta at the 4^th^ to 8^th^ thoracic vertebrae levels were harvested from male mice administered either plain drinking water that was either plain or containing BAPN (0.5% wt/vol) for 7 days (n=20 per group). Periaortic tissues and endothelial cells were removed. Ascending aortas displaying intramural hemorrhage were excluded. Aortic samples not displaying overt pathologies were used for the transcriptomic analysis. Four aortic samples of each group were pooled as one sample for RNA sequencing. The pooled samples were then incubated with RNAlater solution (#AM7020, Invitrogen) overnight. Subsequently, mRNA was extracted using RNeasy Fibrous Tissue Mini kits (#74704, Qiagen) and shipped to Novogene (CA) for mRNA sequencing (n=5 biological replicates per group).

Sequencing library was generated from total mRNA (1 µg) using NEBNext UltraTM RNA Library Prep Kits for Illumina (New England BioLabs, MA). cDNA libraries were sequenced by a NovaSeq 6000 (Illumina) in a paired-end fashion to reach more than 1,500,000 reads. FASTQ sequence data were mapped to mouse genome mm10 using STAR (v2.5, mismatch=2) and quantified using HTSeq (v0.6.1, -m union). For quality control, we removed the following reads: containing > 10% of undetermined bases, Q-score < 5.

### Western Blot Analyses

Aortic tissues were homogenized in Cell Lysis buffer (9803, Cell Signaling Technology) with a protease inhibitor cocktail (P8340, Sigma-Aldrich) using Kimble Kontes disposable Pellet Pestles (Z359971, DWK Life Science LLC.). Protein lysate from HT-1080 cells incubated with human transforming growth factor (TGF)β3 for 30 minutes was used as a positive control, respectively (12052, Cell Signaling Technology). Protein concentrations were determined using a DC assay kit (5000111, Bio-Rad). Equal masses of protein per sample (15 µg) were resolved by SDS-PAGE (10% wt/vol) and transferred to PVDF membranes electrophoretically. After blocking, antibodies against the following proteins were used to probe membranes: p-SMAD2 (0.1 µg/mL, 3108S, Cell Signaling Technology), SMAD2 (0.1 µg/mL, 5339S, Cell Signaling Technology), p-ERK (0.1 µg/mL, 9101S, Cell Signaling Technology), ERK (0.1 µg/mL, 9102S, Cell Signaling Technology), and β-actin (0.3 µg/mL, A5441, Sigma-Aldrich). Membranes were incubated with either goat anti-rabbit (1.0 µg/mL, PI-1000, Vector Laboratories) or goat anti-mouse secondary antibodies (0.3 µg/mL, A2554, Sigma-Aldrich). Immune complexes were visualized by chemiluminescence (34080, Thermo Fisher Scientific) using a ChemiDoc (12003154, BioRad) and quantified using Image Lab software (v6.0.0, BioRad).

### Gelatin Zymography

Aortic samples were homogenized in Cell Lysis buffer and aortic protein (10 µg) was dissolved in zymogram sample buffer (25mM Tris pH6.8, 10% glycerol, 1% SDS, 0.02% bromophenol blue). Subsequently, samples were loaded onto zymography gels (ZY00102BOX, Thermo Fisher Scientific) and run with SDS running buffer for 90 minutes (LC2675, Thermo Fisher Scientific). Gels were renatured in Zymogram Renaturing Buffer (LC2670, Thermo Fisher Scientific) for 30 minutes at room temperature and incubated with Zymogram Developing Buffer (LC2671, Thermo Fisher Scientific) for 30 minutes at 37°C. Gels were incubated further with fresh developing buffer for 72 hours at 37°C. Then, gels were washed with dH_2_O and stained with SimplyBlue SafeStain (LC6060, Thermo Fisher Scientific) for 20 minutes. A ChemiDoc system was used to acquire images of the stained gels.

### Statistical analyses

To compare survival rates within a single factor, Kaplan-Meier Survival curves were generated and a Log-Rank test was used to evaluate the effects of the factor on the survival rates followed by a Holm-Sidak pairwise multiple comparison test (SigmaPlot 15; SYSTAT Software Inc.). Incidence based on aortic rupture location was analyzed by Chi-square in male mice and by Fisher-exact test in female mice. For continuous variables measured after euthanasia, the assumption of normality was examined using QQ-plot and Shapiro-Wilk test. When the assumption was not satisfied, a Box-Cox transformation was used to achieve normality. The homogeneous group variance assumption was assessed by Levene’s test. In studies including both sexes, two-way ANOVA was used. When heteroscedasticity was present, inverse variance weights were incorporated into the two-way ANOVA. Mixed effects model with random intercept and slope of time (week) was fitted to log-transformed ultrasound data implemented by the nlme R package (v3.1). As post hoc tests, contrasts were defined and tested using the glht function in the multcomp R package (v1.4-25)^21^ with the Bonferroni correction. P < 0.05 was considered statistically significant.

Contractility data were analyzed by fitting a sigmoid function using “SSlogis” function in “nlme” R package (v3.1) with three parameters and one dummy variable representing the BAPN group. We added random effects of the parameters and tested the interactions between each of the parameters and the dummy variable to examine whether the sigmoid curves in the control group are equivalent to the BAPN group.

Transcriptome data were analyzed using “edgeR” (v3.36.0) and “clusterProfiler” (v4.2.2) R packages.^22, 23^ Pseudogenes, predicted, and mitochondrial genes were excluded. Principal component analysis was performed using “prcomp” function. False discovery rate (FDR)-adjusted P values were calculated using “p.adjust” function in R. Heatmaps were generated using “pheatmap” R package. Read count data used for the data presented in this manuscript were in the Supplemental Excel File.

### Data availability

The numerical data for figures is provided in a Supplemental Excel File. RNAseq data are available at the Gene Expression Omnibus (GSE241968). The data that support the findings of this study are also available from the corresponding author upon reasonable request.

## RESULTS

### Dose-response curve of BAPN administration in young male C57BL/6J mice

There is a paucity of data on the influence of BAPN dosage provided in drinking water on development of aortopathies.^6^ Therefore, we first performed a dose-response curve for BAPN in 3-week-old male C57BL/6J mice (Figure 1A). BAPN in drinking water at concentrations of 0.1%, 0.3%, or 0.5% (wt/vol) was administered for 12 weeks (N=10/concentration). No death occurred in mice drinking 0.1 % BAPN, but 6 of 10 mice and 8 of 10 mice died of aortic rupture in the groups drinking 0.3% and 0.5% of BAPN, respectively (Figure 1A). In mice that survived for 12 weeks, 2 mice (20%) in BAPN 0.1% group and 2 of the 4 survived mice (50%) in BAPN 0.3% group had apparent dilatations in the ascending aortic region, but no overt pathologies were noted in the descending aortic region of either group. In addition to the high aortic rupture rate, the two mice that survived in BAPN 0.5% group had profound dilatation of the ascending aortic region, and one mouse also had pathologies in the descending thoracic region. Therefore, in subsequent studies, BAPN was provided in drinking water at a concentration of 0.5% wt/vol.

**Figure 1.**
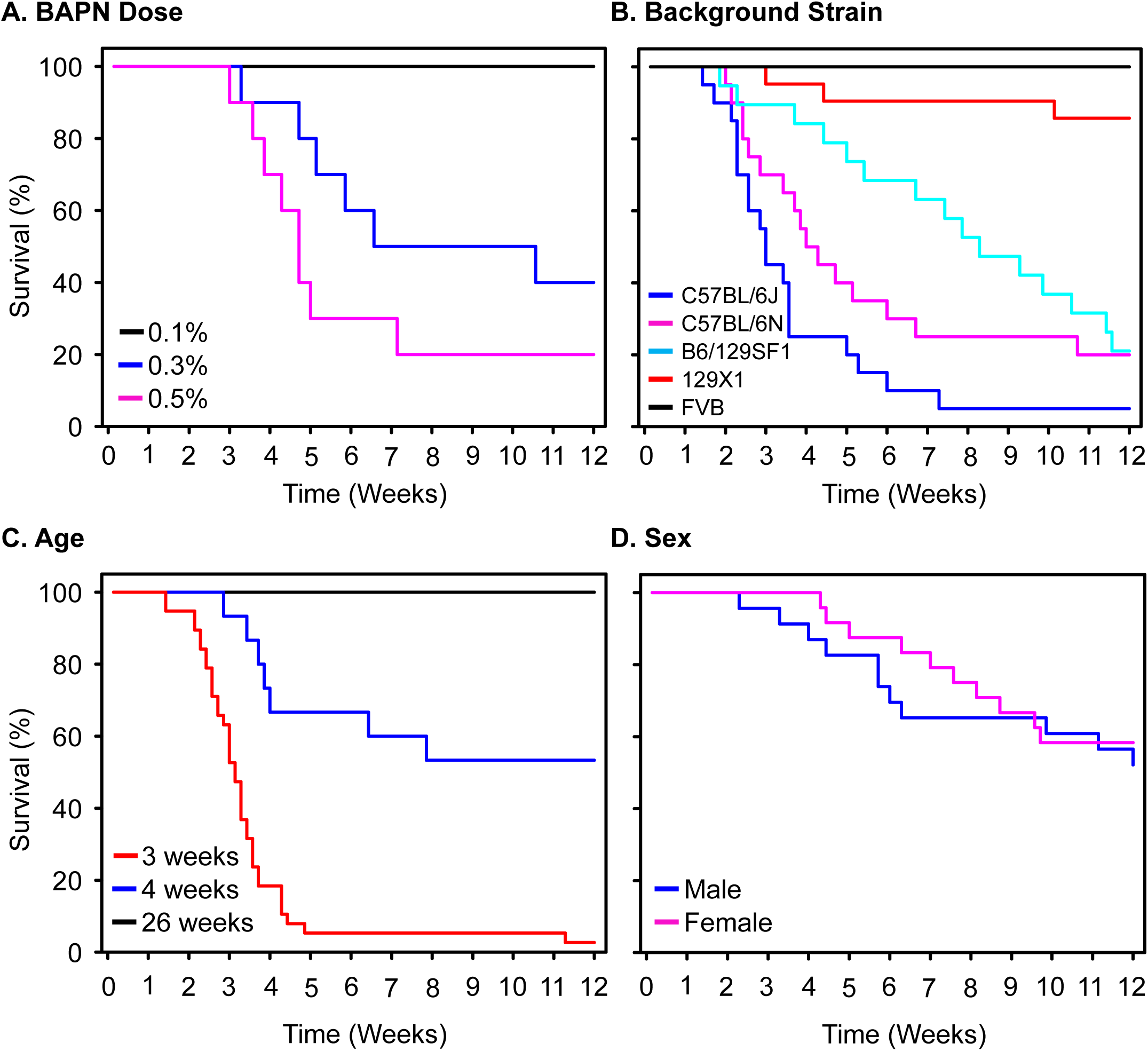
Survival curves of BAPN-induced aortic rupture. Survival curves of **(A)** male C57BL/6J mice administered BAPN 0.1%, 0.3%, or 0.5% wt/vol (N=10/concentration) for 12 weeks, **(B)** five mouse strains administered BAPN 0.5% (N=19-21/mouse strain, male), **(C)** male C57BL/6J mice started BAPN 0.5% at 3 (N=38), 4 (N=15), or 26 (N=15) weeks of age, and **(D)** male (N=24) and female (N=23) C57BL/6J mice administered BAPN 0.5%. P values were determined by Log-Rank analysis. P=0.008 for BAPN 0.1% vs 0.3% and P<0.001 for BAPN 0.1% vs 0.5% (A); P values for (B) are detailed in Table S1; P=0.003 for 4- and 26-week comparison, and P<0.001 for 3- vs 4-week comparison and 3- vs 26-week comparison (C). BAPN administration was started at 3 weeks of age in (A) and (B), and 4-weeks of age in (D).

### Impact of mouse strains on the susceptibility to BAPN-induced aortopathies

A previous study reported that 3-week-old male FVB mice had much lower susceptibility to aortopathies, compared to age-matched male C57BL/6 (substrain not described) mice.^12^ We therefore compared BAPN-induced aortopathies in 5 mouse strains (C57BL/6J, C57BL/6N, B6/129SF1, 129X1, and FVB) that are commonly used for genetic manipulations. Male mice (N=19-21/mouse strain) at 3 weeks of age were administered BAPN (0.5% wt/vol) in drinking water. Among the 5 strains, C57BL/6J mice (19 of 20 mice) and C57BL/6N (16 of 20 mice) had comparable death rate due to aortic rupture; B6/129SF1 mice (15 of 19 mice) had delayed aortic rupture, compared to C57BL/6J. Three of 21 mice with 129X1 strain died of aortic rupture, but no death occurred in FVB mice (Figure 1B and Table S1). Apparent pathologies were noted in C57BL/6J, C57BL/6N, and B6/129SF1 mice, and less severe pathologies were observed in 129X1 mice, while only 2 FVB mice had modest pathologies in either the ascending or descending thoracic region (Table S2, Figures S1 and S2). Based on the C57BL/6J and C57BL/6N mouse strains having the most severe aortopathies, and our research projects have predominantly used mice with C57BL/6J background, subsequent studies used this substrain.

### Effects of age and sex on BAPN-induced aortopathies

To determine the impact of age on BAPN-induced aortopathies, male C57BL/6J mice were administered BAPN (0.5% wt/vol) at 3, 4, or 26 weeks of age (Figure 1C). No death or overt pathologies were found in mice started BAPN at 26 weeks of age. In contrast, death due to aortic rupture was noted after 10-14 days of BAPN administration in mice that were initiated BAPN at 3 or 4 weeks of age, and the death rate was much higher if BAPN was started at 3 weeks of age, compared to 4 weeks of age.

No studies have previously compared BAPN-induced aortopathies between male and female mice side-by-side within an experiment. To determine whether BAPN-induced aortopathies exhibit sex differences, 4-week-old male and female C57BL/6J mice were administered BAPN (0.5% wt/vol). Although male mice succumbed to aortic rupture earlier and in greater numbers than females during the first 4 weeks of BAPN administration, no difference of aortic rupture rate was found between sexes after prolonged administration of BAPN for up to 12 weeks (Figure 1D).

In mice that died before termination, necropsy revealed that deaths were attributed mainly to aortic rupture that was restricted or initiated in the descending thoracic region (Figure 2A and 2B). Among 78 male mice given BAPN at 3-4 weeks of age, 13% (N=10) died of ascending aortic rupture, 87% (N=68) died of descending aortic rupture that was either restricted to the descending thoracic region or involved both the descending thoracic region and the abdominal region. Comparably, female mice (N=21) had 14% and 86% of aortic rupture in the two aortic regions, respectively.

**Figure 2.**
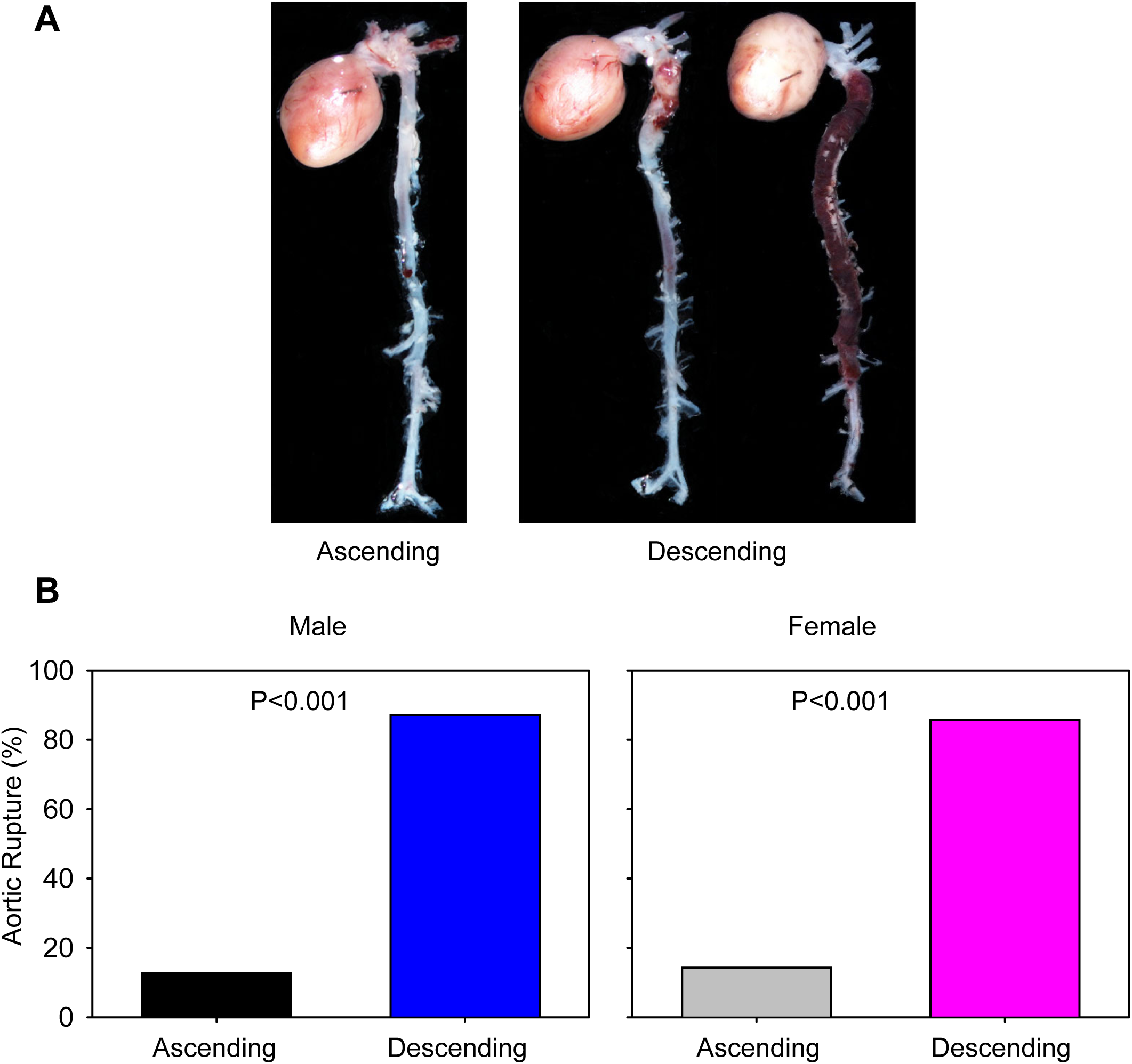
Locations of BAPN-induced aortic rupture in young C57BL/6J mice. BAPN (0.5% wt/vol) was administered in drinking water for 12 weeks. Necropsy was performed for mice that died during BAPN administration within 12 weeks. **(A)** Examples of ex vivo images showing ascending or descending aortic rupture. **(B)** Death rate attributed to aortic rupture in the ascending versus descending aortic regions of male and female mice. Necropsy was performed on 78 male mice and 21 female mice. Incidence based on aortic rupture location was analyzed by Chi-square in male mice (P<0.001), and by Fisher-exact test in female mice (P<0.001).

### BAPN promoted aortic dilatation in the ascending and descending thoracic regions, but not abdominal aortic region

To monitor luminal dilatations in the ascending aortic region, ultrasound was performed in both male and female mice to determine luminal diameters (1) between mice administered control (drinking water only) and BAPN for 12 weeks, and (2) between mice started BAPN at 4 weeks and 26 weeks of age. In mice started BAPN at 4 weeks of age, luminal diameters were significantly larger in both male and female mice administered BAPN for 12 weeks, and no differences of luminal diameters of ascending aortas were noted between young male and female mice administered BAPN (Figures 3A and 3B). Administration of BAPN led to larger luminal diameters in young mice than in mature mice within each sex; however, in mice that were started with BAPN at 26 weeks of age, no luminal dilatations were observed in either sex (Figures 3C and 3D).

**Figure 3.**
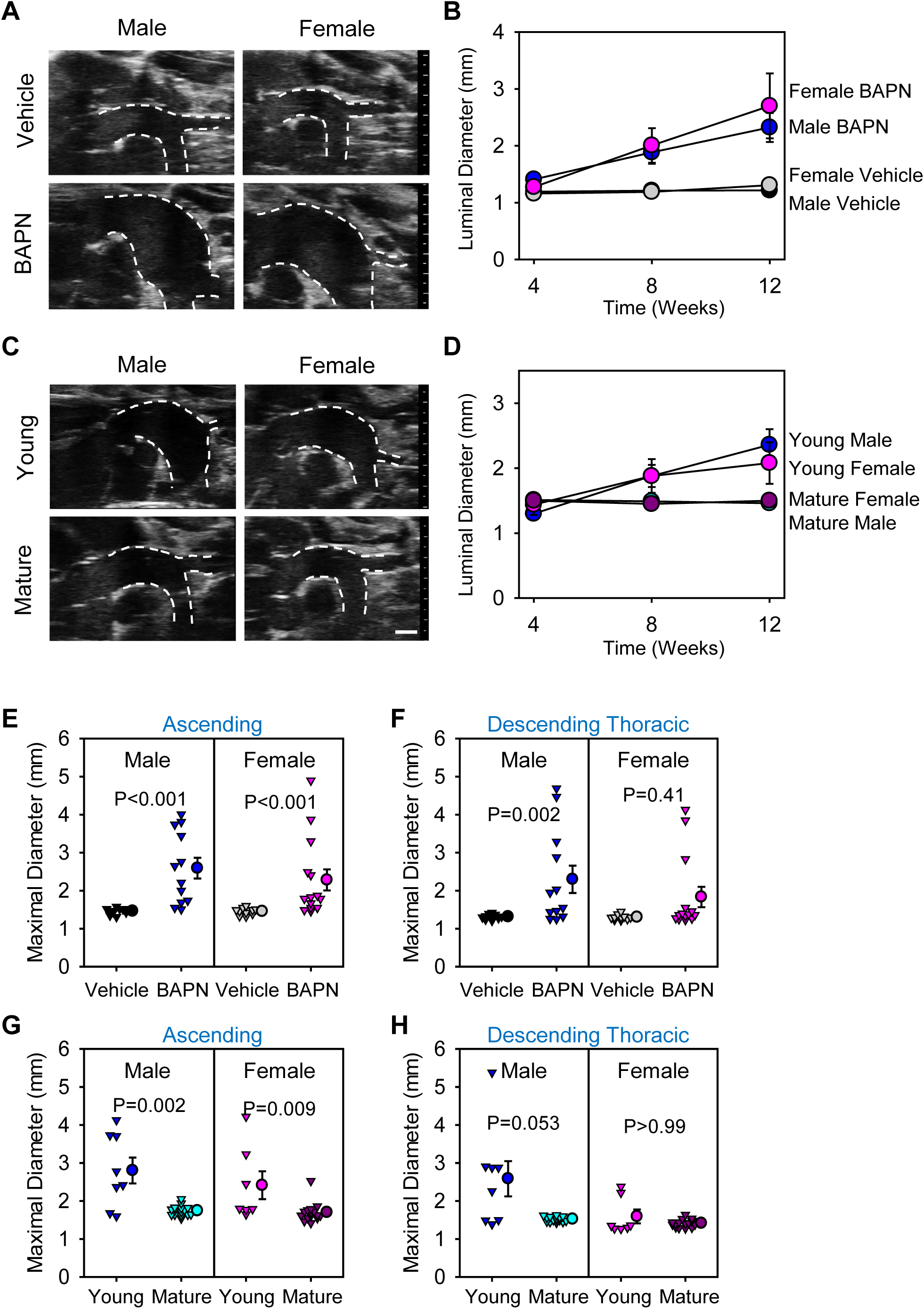
BAPN induced dilatation of the ascending and descending thoracic regions in young mice. Representative ultrasound images and luminal diameters of ascending aortas in young (4-week-old) C57BL/6J mice administered vehicle (drinking water only) versus BAPN **(A and B)** and in young (4-week-old) versus mature (26-week-old) mice administered BAPN **(C and D)**. BAPN versus vehicle after 12 weeks of BAPN administration in male and female mice: P<0.001 in both sexes, and the difference in changes over time between male and female mice of BAPN administration: P>0.99. **(B)**. Young versus Mature mice within males and females: P<0.001 and P=0.015, respectively, and the difference in changes over time between young male and female mice: P=0.16. **(D)**. In situ measurements of maximal diameters of ascending/arch regions **(E)** and descending thoracic regions **(F)** in 4-week-old C57BL/6J mice administered vehicle versus BAPN for 12 weeks. In situ measurements of maximal diameters of ascending/arch regions **(G)** and descending thoracic regions **(H)** in 4-week-old (Young) versus 26-week-old (Mature) C57BL/6J mice administered BAPN for 12 weeks. P=0.56 between male and female mice administered BAPN **(E)**, P=0.096 between male and female mice administered BAPN **(F)**, P=0.93 between male and female young mice administered BAPN **(G)**, and P=0.086 between male and female young mice administered BAPN **(H)**. P values in B and D were determined using the mixed effects model with post hoc contrast tests. Data in E-H were analyzed using two-way ANOVA with post hoc contrast tests within each sex.

In mice surviving 12 weeks of BAPN administration, maximal external diameters were measured in the ascending and descending thoracic regions using in situ images (Figure 3E-H), and abdominal aortic diameters were measured using ex vivo images (Figure S3). Aortic expansions were profound in the ascending and descending thoracic aortic regions in male young mice administered BAPN, compared to the age-matched vehicle group (Figure 3E-F), and compared to male mature mice administered BAPN (Figure 3G-H). Increases of diameters were only statistically significant in the ascending, but not descending thoracic region in young female mice when compared to either age-matched vehicle group (Figure 3E-F) or mature mice administered BAPN (Figure 3G-H). No overt dilatations were found in the abdominal aortic region of young male or female mice administered BAPN, compared to their age-matched vehicle group or sex-matched mature mice administered BAPN.

Based on these results, subsequent studies for histological characterization focused on the ascending and descending thoracic aortic regions of male C57BL/6J mice administered BAPN (0.5% wt/vol) in drinking water that was provided at ∼4 weeks of age.

### Temporal characteristics of BAPN-induced thoracic aortopathies

To determine temporal evolution of BAPN-induced thoracic aortopathies, male mice were terminated at different intervals of BAPN administration (started at ∼4 weeks of age): baseline, 1, 2, 3, or 4 weeks (Figure 4A). Consistent with the survival curve that no mice died during the first week, discernable pathology was not observed in the aorta after 1 week of BAPN administration. Overt pathology was first noted after 8-9 days of BAPN administration. Among 15 mice administered BAPN for 9, 12, 14, 17, or 21 days, gross pathology was observed more frequently in the ascending and arch region (10 of 15 mice; 67%) compared to the descending thoracic region (3 of 15 mice; 20%) of the survived mice. Two of 15 mice (13%) had pathologies at both the ascending/arch region and the descending thoracic aortic region. In mice with acute aortic dissections, thrombus between elastic layers that also led to a false lumen was noted by H&E staining in the ascending/arch or descending thoracic region (Figure 4B).

**Figure 4.**
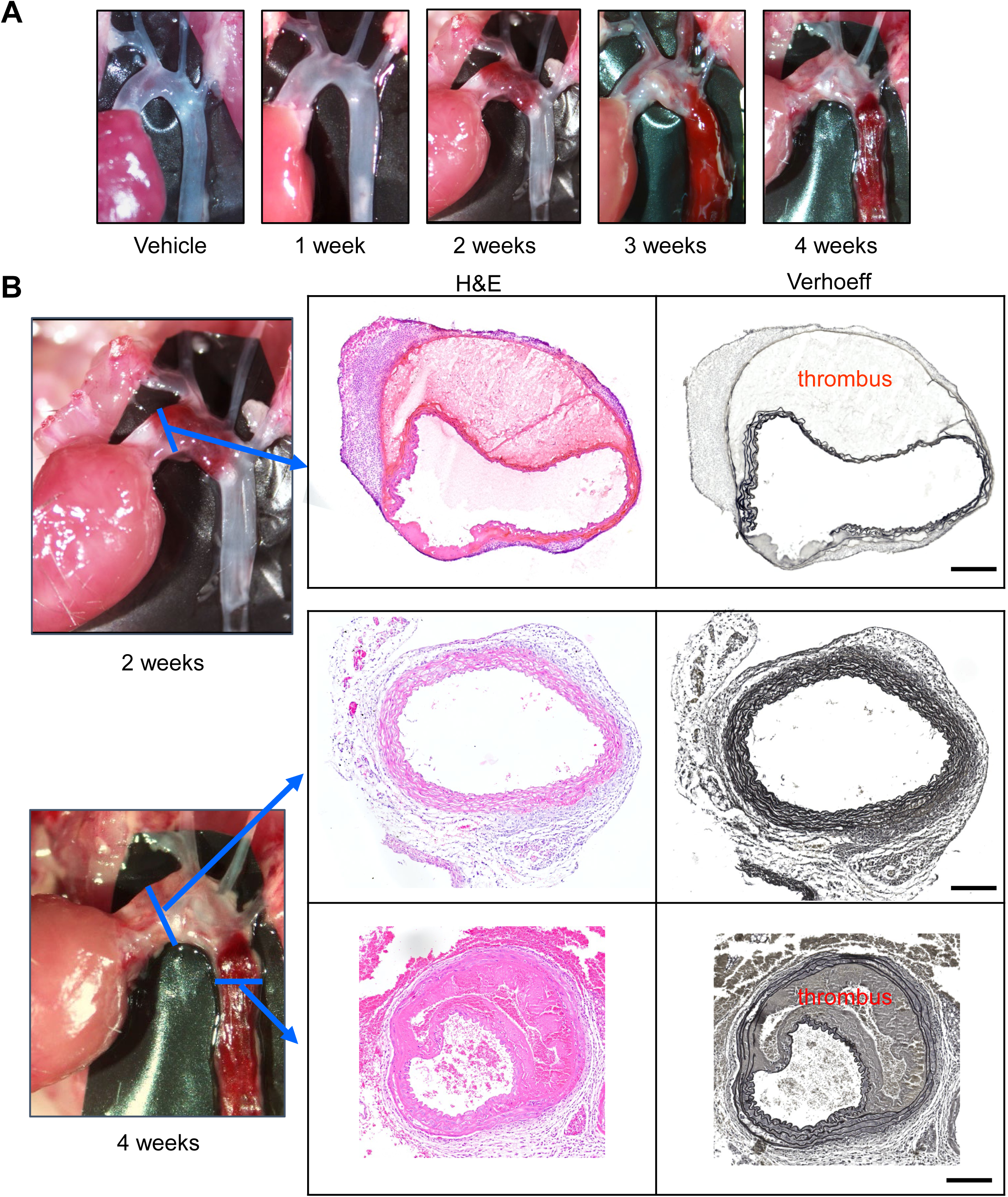
Temporal characteristics of BAPN-induced thoracic aortopathies in young male C57BL/6J mice. BAPN was administered in drinking water for 1, 2, 3, or 4 weeks in 3-4-week-old male C57BL/6J mice. **(A)** Examples of in situ images of thoracic aortas. **(B)** Hematoxylin-eosin (H&E) and Verhoff staining images of ascending aortic dissection and descending thoracic dissection. Scale bar is 200 µm.

### Prolonged administration of BAPN led to profound vascular remodeling in the dissected descending thoracic aortic region

As shown in Figure 2, in mice surviving 12 weeks of BAPN administration, aortic dissection in the descending thoracic aorta became more frequent, while aortic pathology was predominant with dilatations in the ascending/arch region. Histological staining (H&E, Verhoeff, or Movat) and immunostaining of α-smooth muscle actin (α-SMA) and macrophage (CD68) were performed in tissue sections from the ascending and descending thoracic aortas of mice after 12 weeks of BAPN administration. Aortic pathologies in the ascending/arch region were characterized by profound medial thickening or thinning and elastic fiber disruption (Figures 5 and Figure S4-6). Development of false lumens was infrequent in the ascending aortic region, whereas thickening and remodeling of false lumen were noticed frequently in the descending thoracic aorta (Figure 5 and Figures S2 and S4). In the aortic wall of the descending thoracic aorta circumscribed to the true lumen, elastic fibers in the aortic media were disrupted and collagen fibers were deposited in the adventitia, and elastic fiber fragmentation and collagen deposition were more profound in the vascular wall of the false lumen (Figure S4). Immunostaining revealed that the vascular wall of the false lumen was covered by α-SMA positive cells, accompanied by CD31 positive cells in the intima and COL1A1 and CD68 positive cell accumulation in the adventitia (Figure 5, Figure S5). There was no obvious accumulation of cells expressing either CD3 or CD19 (Figure S5).

**Figure 5.**
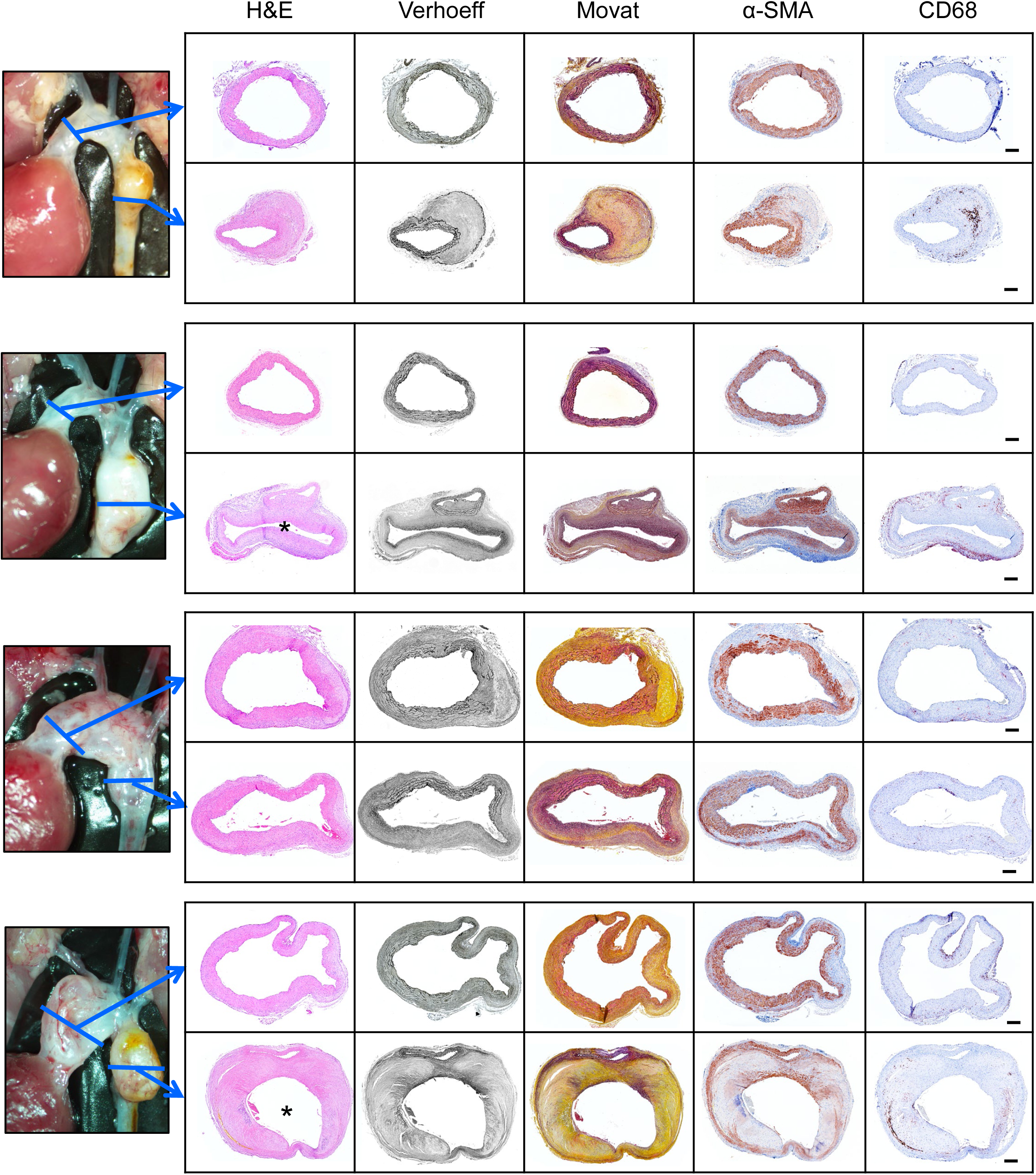
Chronic administration of BAPN led to profound vascular remodeling following dissection in the descending thoracic region. Representative images of hematoxylin-eosin (H&E), Verhoeff iron hematoxylin, Movat’s pentachrome, α-SMA (smooth muscle actin), and CD68 staining in ascending and descending aortas of mice with BAPN administration for 12 weeks. The approximate location of the tissue section from the ascending and descending region is indicated by the blue line. * indicates false lumen. Scale bar is 200 µm.

### BAPN led to similar changes of contractility and transcriptomics between the ascending and descending thoracic aortic regions in a pre-pathological phase

It has been proposed that compromise of aortic contractile units is a precipitating factor in the development of aortopathies.^24–26^ Therefore, the contractile properties were determined in the ascending and descending aortic regions after 1 week of either vehicle or BAPN administration prior to aortopathy formation (Figure S7). Concentration-based contractility curves were developed based on the incubation of aortic rings with 5-hydroxytryptamine ex vivo. Compared to vehicle-administered mice, the maximum contractile force in response to 5-hydroxytryptamine was decreased in both regions of BAPN-administered mice (Figure S7).

To investigate the potential molecular basis of the regional specificity of BAPN-induced thoracic aortopathy, RNA sequencing was performed using ascending and descending aortic samples harvested following 7 days of BAPN administration after confirming no overt pathological changes in the aortic wall. Principal component analysis revealed that BAPN administration altered aortic transcriptomes in a region-specific manner (Figure 6A). Two-way ANOVA for the interaction between administration and sex only identified 12 genes (Figure 6B). However, there were no obvious differences between the two aortic regions in response to BAPN in cell proliferation markers, smooth muscle cell markers, or extracellular components that have been reported to link to the pathophysiology of thoracic aortopathy (Figure 6C). Since there is compelling evidence indicating the contribution of TGFβ signaling and matrix metalloproteinase (MMP) activity to thoracic aortopathy formation,^27^ aortic SMAD2 and ERK phosphorylation and MMP2/ MMP9 activities were assessed. Western blot analysis demonstrated that neither pSMAD2 nor pERK was increased in the aortas of BAPN-administered mice prior to development of overt pathology, regardless of the aortic region (Figure 6D). Of note, while MMP9 was minimally abundant in either aortic region, the active/latent MMP2 ratio in the descending thoracic region was higher than that in the ascending aortic region (Figure 6E).

**Figure 6.**
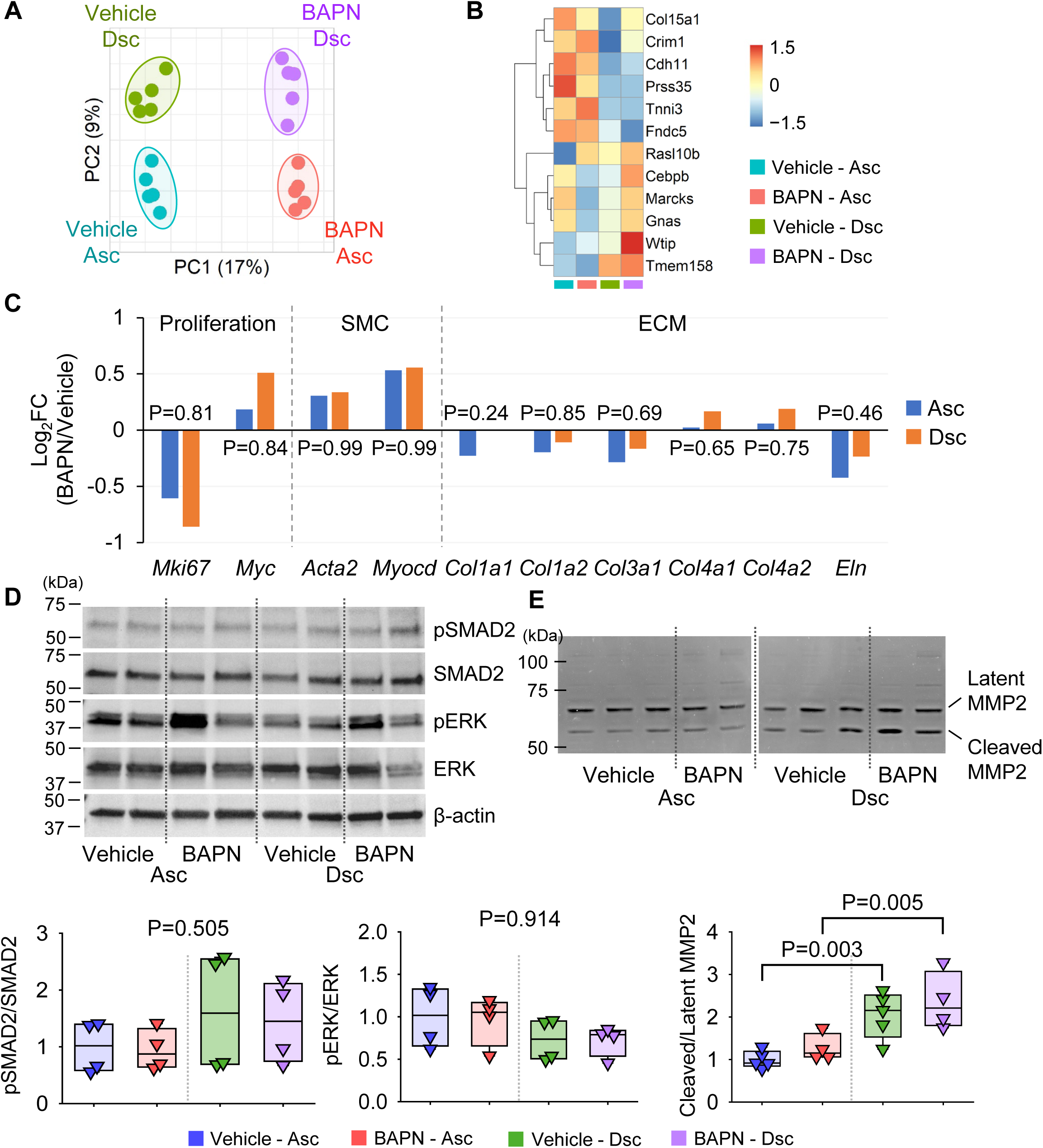
Transcriptomic alteration by BAPN administration in the ascending and descending thoracic aortas. **(A)** Principle component analysis for unfiltered transcriptomes of ascending and proximal descending thoracic aortas harvested from 4-week-old male C57BL/6J mice administered either vehicle or BAPN for 7 days. **(B)** Heatmap depicting z-scored normalized read counts of differentially expressed genes (DEGs) from the interaction analysis between sexes and administration in two-way ANOVA. **(C)** Bar plots for DEGs related to proliferation, smooth muscle cell contraction, the extracellular maturation (ECM). N=5 biological replicates per group. **(D)** Western blot and **(E)** gelatin zymography and their quantifications. N=4-5/group and P values were calculated by Kruskal-Wallis test (pSMAD2) or two-way ANOVA followed by Holm-Sidak method (pERK and MMP2).

## DISCUSSION

Recently there has been greatly increased interest in BAPN-induced aortopathies, but there is a lack of knowledge on many fundamental issues regarding this mouse model. The present study reports several new findings including (1) BAPN-induced aortopathies have a mouse strain-specific susceptibility, with C57BL/6J and C57BL/6N being the most susceptible mouse strains; (2) the onset of BAPN-induced aortic rupture is delayed in females within the first 4 weeks of BAPN administration; however, BAPN-induced aortopathies do not show sex differences during its chronic phase (up to 12 weeks of BAPN administration); (3) aortic rupture rate was higher in mice started BAPN administration at 3 weeks of age than at 4 weeks of age; (4) BAPN-induced aortopathies are highly heterogeneous; (5) BAPN-induced aortic dissections and rupture are more frequent in the descending aortic region; (6) a striking feature of the BAPN-induced pathologies was the profound differences between the ascending and descending regions of the aorta; chronic administration (up to 12 weeks) of BAPN administration led to remodeling of the vascular wall of the false lumen in the descending thoracic region.

LOX and LOX-like proteins (LOX-L1 to L4) are responsible for crosslinking of elastic fibers and collagen, with all being present in the aorta.^28–32^ Mutations of LOX with reduced enzymatic activity are linked to the aggressive development of human thoracic aortopathy in the aortic root and ascending aortic region.^29, 31, 33^ Mice with deletion or missense mutations of LOX create live births of homozygous mice that die shortly after birth with hemorrhage in the aorta and other loci.^29, 31^ The aortas of mice with missense mutations had ascending and arch aortic aneurysms and pronounced tortuosity throughout the descending aorta.^29^ Therefore, constitutive deletion of LOX activity profoundly affects aortic development. A study reported that LOXL4 deletion had no effects on AngII-induced aortic aneurysms in mice.^34^ Currently there are no studies regarding aortic pathologies on LOXL1 to L3 of this family.

In contrast to the limited genetic studies,^29, 33^ the current study determined the effect of pharmacological inhibition of LOX and LOX-like proteins in the postnatal phase using BAPN. BAPN is considered to be an irreversible inhibitor of LOX and LOX-like proteins, although there is an inconsistent literature on the specificity of the drug to inhibit these five proteins.^35, 36^ BAPN has been administered to mice using several different approaches including in diet,^37^ in osmotic pumps,^9, 10^ through gastric tube,^38^ or intraperitoneal injection,^39^ but the most frequently used approach was given in drinking water.^6, 12, 16, 19, 40, 41^ The current study also provided BAPN in drinking water with determination of a dose-dependent effect.

Using the maximal dose (BAPN 0.5% wt/vol) from the initial studies in male C57BL/6J mice, we subsequently studied the effects of this dose side-by-side on C57BL/6J and four other strains that were selected based on their common use as the background strains in aortopathy studies. In agreement with a previous study,^12^ we found this dose of BAPN failed to cause aortic rupture in FVB mice that also had no or modest aortic pathologies after 12 weeks of BAPN administration. A recent study reported that C57BL/6J mouse strain is more susceptible to AngII-induced abdominal aortic aneurysm than C57BL/6N mouse strain.^42^ For mice administered BAPN, there were no discernable differences between the rate of death due to aortic rupture in C57BL/6J and C57BL/6N mice. This finding will have relevance to many studies since many genetically modified mice originate from the C57BL/6N strain. Stem cells from B6/129SF1 and 129X1 strains are also frequently used in creating genetically manipulated mice. Their prevalence of death attributed to aortic rupture was intermediate between the 5 strains examined. Although the mechanisms by which different mouse strains have different susceptibility to BAPN-induced aortopathies are undefined, the present study has demonstrated the importance of providing precise descriptions of mouse strain in studies using BAPN.

Many forms of experimental aortopathies have a strong sexual dimorphism.^43–46^ Previous studies with BAPN administration have either used male mice or not provided the sex information. Overall, there has not been direct comparisons between male and female mice. In the current studies, there appeared to be a sexually dimorphic response with male mice succumbing to aortic rupture earlier than female mice during the first 4 weeks of BAPN administration. However, during 12 weeks of BAPN administration, female mice had equivalent death attributed to aortic rupture as male mice, and maximal diameters in the ascending aortic regions were not significantly different between the two sexes. Hence, there was a delay in the initiation of aortopathies in female mice, of which the mechanisms will need further investigation. In contrast, a previous study using BAPN administration with topical elastase application reported that female mice have accelerated aneurysm growth, enhanced dilatation, and higher mortality compared to their male counterparts.^47^ Multiple factors may contribute to the differences between the previous and our studies, warranting further investigation to elucidate the underlying mediating mechanisms.

Previous studies used 3-week-old (prior to sexual maturation) C57BL/6 mice when BAPN alone was administered.^6, 12^ The current study demonstrated that initiation of BAPN at 3 or 4 weeks of age led to high aortic rupture rate and profound aortic pathologies, whereas aortic rupture or pathologies were not detected when BAPN administration was initiated at 26 weeks of age. Our findings in mature mice were also supported by a previous study showing that 9-week-old male C57BL/6J mice did not develop aortopathies when BAPN alone was administered.^9^ BAPN is an irreversible inhibitor of LOX and LOX-like proteins that is presumed to not affect already cross-linked elastic fibers.^6, 48^ This age-dependency of BAPN-induced aortopathies may reflect that elastic fibers are actively crosslinked for their maturation when mice are 3-4 weeks old, while elastic fibers in the aortic wall have already been mature in 26-week-old mice, which do not require crosslinking of elastic fibers unless pathological insults lead to disruption of the already formed elastic fibers.^9, 48–50^ It is also worth noting that aortic rupture rate was higher in mice started BAPN administration at 3 weeks of age than at 4 weeks of age. This finding further confirmed the age-sensitive effects of BAPN in aortopathy formation.^9^ Loss-of-function mutations in the *LOX* gene have been identified in humans.^29, 33^ These patients exhibit aortic pathologies, such as thoracic aortic aneurysms and dissections, from an early age. In addition, patients with connective tissue diseases, such as Marfan syndrome, Ehlers-Danlos syndrome, or Turner syndrome, often encounter aortic manifestations with disruption of elastic fibers. These complications typically initiate at an early age, reaching its peak between 30 to 40 years of age in patients with these syndromes.^51–53^ Therefore, administration of BAPN to young mice replicates aortic manifestations observed in patients with genetic disorders. Under normal development condition, mRNA abundance of tropoelastin, the precursor of elastin, remains high during the early postnatal phase, but reduced strikingly 4 weeks after birth.^54, 55^ While the abundance of tropoelastin mRNA may be associated with BAPN-induced aortopathies, there is limited evidence of the specific changes to elastic fibers during the postnatal phase in normal conditions and under a diseased condition.

For mice surviving 12 weeks of BAPN administration, aortopathies occurred in both ascending and descending thoracic aortic regions, but not the abdominal aortic region. For mice that died of aortic rupture, blood was detected occasionally in the abdominal cavity at necropsy, but all these mice also had dissections originating in the descending thoracic region. For mice survived to the endpoint, BAPN did not cause pathology in the abdominal aorta without involvement of the descending thoracic region. These data support the notion that dissections are initiated from the descending thoracic, not the abdominal aortic, region. In mice, ascending and arch region contains 8-12 elastic layers, descending thoracic region contains 6-8 elastic layers, whereas abdominal aorta only contains 3-5 elastic layers.^56^ It is possible that elastic layers are almost formed and mature in the abdominal aorta at ∼3-4 weeks of age, but the thoracic aortic region with more elastic layers requires more time for maturation of all elastic layers.

Although heterogeneity was noted in both the ascending/arch and descending thoracic aortic regions during chronic administration of BAPN, descending thoracic pathologies showed a wider range of variations in the gross appearance, while the ascending aorta predominantly had luminal dilatation. Also, ascending aortic dilatations were detected earlier than in the descending thoracic region. These data suggest that the ascending aorta is more prone to luminal dilatations, whereas the descending thoracic aorta is more susceptible to medial dissections and formation of false lumens. Several mechanisms have been proposed as a contributing factor to this regional specificity, including hemodynamic effects of the blood flow, anatomic differences, and embryonic origins between the two aortic regions. To explore the distinct pathogenesis of these two aortic regions, we compared contractility and transcriptomics between the ascending and descending thoracic aortic regions in a pre-pathological phase, namely that BAPN was administered for 1 week. Surprisingly, this short-term of BAPN administration led to comparable changes of contractility and transcriptomics between the two regions. Aortic SMAD2 and ERK phosphorylation were also comparable between the regions. However, aortic MMP2 activity was significantly higher in the descending region compared to the ascending region of BAPN-administered mice in the prepathological phase. Since MMP2 is involved in the disruption of aortic ECM,^27^ the regional difference in aortic MMP2 activity may contribute to the regional heterogeneity of BAPN-induced thoracic aortopathy.

In the present study, aortic dissections were observed more frequently in the proximal descending thoracic aorta adjacent to the left subclavian artery. a distinct boundary of BAPN-induced aortic dissection was observed consistently just distal to the left subclavian artery. This location corresponds with the transition of embryonic origins of SMCs.^57–59^ The embryonic origin of SMCs in the aortic arch is the cardiac neural crest,^58, 59^ whereas that of the descending thoracic aorta is the somites.^60^ Therefore, the boundary of BAPN-induced aortic dissection aligns with the interface of embryonic origins between the cardiac neural crest and somite. Unfortunately, there are currently no promoters that drive authentic *Cre* expression in somites-derived cells to investigate the impact of the interface of embryonic origins of SMCs on development and progression of BAPN-induced aortopathies.

The present study compared contractile responses between ascending and descending thoracic aortic regions of BAPN-administered mice. The measurements revealed that biomechanics, as determined by active contraction of smooth muscle cells in response to 5-HT as an agonist, did not differ between the two regions at the prepathological phase in BAPN-administered mice. However, the thoracic aorta, particularly of the ascending region, exhibited extension, distension, and torsion with each cardiac cycle.^61–63^ Therefore, biaxial measurements are needed to further understand the biomechanics for active and passive constitutive relations between the ascending and descending thoracic regions in BAPN-induced thoracic aortopathy.

In conclusion, this study demonstrates that BAPN induces distinct pathologies in the ascending/arch and descending thoracic aortic regions of young mice. Future studies will focus on defining the molecular mechanisms driving the distinct pathological features between the two thoracic aortic regions.

## Acknowledgment

Histological and immunohistochemical images were acquired using Zeiss Axioscan Z1 or 7 in the Light Microscopy Core at the University of Kentucky. Luminal diameters of aortas were measured using Vevo 3100 in the Ultrasound Core at the University of Kentucky.

## Author Contributions

MKF, HS, SI, DAH, NA, C-LL, NZ, DBG, JJM, and HSL performed the experiments and data verification. MKF, HS, SI, YK and HSL analyzed the data. MKF, HS, HSL, and AD drafted and revised the manuscript. HSL and AD designed and supervised the experiments. All authors contributed to manuscript editing and provided the final approval to the manuscript.

## Ethics Approval

All research procedures related to mice were approved by the University of Kentucky Institutional Animal Care and Use Committee.

## Highlights

1. Administration of BAPN to 3-4-week-old C57BL/6J mice leads to a high incidence of descending thoracic aortic dissection and rupture.
2. Pathological features are distinct between the ascending and descending thoracic regions in BAPN-administered young mice.
3. Aortic dissection and rupture rates are not different between male and female young C57BL/6J mice administered BAPN for prolonged intervals.

## REFERENCES

1. Shen YH, LeMaire SA, Webb NR, Cassis LA, Daugherty A and Lu HS. Aortic aneurysms and dissections series. Arterioscler Thromb Vasc Biol. 2020;40:e37–e46.

2. Tsamis A, Krawiec JT and Vorp DA. Elastin and collagen fibre microstructure of the human aorta in ageing and disease: a review. J R Soc Interface. 2013;10:20121004.

3. Heinz A. Elastic fibers during aging and disease. Ageing Res Rev. 2021;66:101255.

4. Yanagisawa H and Wagenseil J. Elastic fibers and biomechanics of the aorta: Insights from mouse studies. Matrix Biol. 2020;85-86:160–172.

5. Schmelzer CEH, Hedtke T and Heinz A. Unique molecular networks: Formation and role of elastin cross-links. IUBMB Life. 2020;72:842–854.

6. Sawada H, Beckner ZA, Ito S, Daugherty A and Lu HS. Beta-Aminopropionitrile-induced aortic aneurysm and dissection in mice. JVS Vasc Sci. 2022;3:64–72.

7. Bachhuber TE and Lalich JJ. Production of dissecting aneurysms in rats fed Lathyrus odoratus. Science. 1954;120:712–713.

8. Lalich JJ, Barnett BD and Bird HR. Production of aortic rupture in turkey poults fed beta-aminopropionitrile. AMA Arch Pathol. 1957;64:643–648.

9. Kanematsu Y, Kanematsu M, Kurihara C, Tsou TL, Nuki Y, Liang EI, Makino H and Hashimoto T. Pharmacologically induced thoracic and abdominal aortic aneurysms in mice. Hypertension. 2010;55:1267–1274.

10. Remus EW, O’Donnell REJ, Rafferty K, Weiss D, Joseph G, Csiszar K, Fong SF and Taylor WR. The role of lysyl oxidase family members in the stabilization of abdominal aortic aneurysms. Am J Physiol Heart Circ Physiol. 2012;303:H1067–1075.

11. Takayanagi T, Crawford KJ, Kobayashi T, Obama T, Tsuji T, Elliott KJ, Hashimoto T, Rizzo V and Eguchi S. Caveolin 1 is critical for abdominal aortic aneurysm formation induced by angiotensin II and inhibition of lysyl oxidase. Clin Sci (Lond). 2014;126:785–794.

12. Ren W, Liu Y, Wang X, Jia L, Piao C, Lan F and Du J. beta-Aminopropionitrile monofumarate induces thoracic aortic dissection in C57BL/6 mice. Sci Rep. 2016;6:28149.

13. Xu K, Xu C, Zhang Y, Qi F, Yu B, Li P, Jia L, Li Y, Xu FJ and Du J. Identification of type IV collagen exposure as a molecular imaging target for early detection of thoracic aortic dissection. Theranostics. 2018;8:437–449.

14. Gao Y, Wang Z, Zhao J, Sun W, Guo J, Yang Z, Tu Y, Yu C, Pan L and Zheng J. Involvement of B cells in the pathophysiology of β-aminopropionitrile-induced thoracic aortic dissection in mice. Exp Anim. 2019;68:331–339.

15. Wang F, Tu Y, Gao Y, Chen H, Liu J and Zheng J. Smooth muscle sirtuin 1 blocks thoracic aortic aneurysm/dissection development in mice. Cardiovasc Drugs Ther. 2020;34:641–650.

16. Zheng HQ, Rong JB, Ye FM, Xu YC, Lu HS and Wang JA. Induction of thoracic aortic dissection: a mini-review of β-aminopropionitrile-related mouse models. J Zhejiang Univ Sci B. 2020;21:603–610.

17. Wortmann M, Arshad M, Peters AS, Hakimi M, Bockler D and Dihlmann S. The C57Bl/6J mouse strain is more susceptible to angiotensin II-induced aortic aneurysm formation than C57Bl/6N. Atherosclerosis. 2020;318:8–13.

18. Sawada H, Chen JZ, Wright BC, Moorleghen JJ, Lu HS and Daugherty A. Ultrasound imaging of the thoracic and abdominal aorta in mice to determine aneurysm dimensions. J Vis Exp. 2019:10.3791/59013.

19. Ohno-Urabe S, Kukida M, Franklin MK, Katsumata Y, Su W, Gong MC, Lu HS, Daugherty A and Sawada H. Authentication of in situ measurements for thoracic aortic aneurysms in mice. Arterioscler Thromb Vasc Biol. 2021;41:2117–2119.

20. Ito S, Lu HS, Daugherty A and Sawada H. Imaging techniques for aortic aneurysms and dissections in mice: comparisons of ex vivo, in situ, and ultrasound approaches. Biomolecules. 2022;12:339.

21. Hothorn T, Bretz F and Westfall P. Simultaneous inference in general parametric models. Biom J. 2008;50:346–363.

22. Robinson MD, McCarthy DJ and Smyth GK. edgeR: a Bioconductor package for differential expression analysis of digital gene expression data. Bioinformatics. 2010;26:139–140.

23. Wu T, Hu E, Xu S, Chen M, Guo P, Dai Z, Feng T, Zhou L, Tang W, Zhan L, Fu X, Liu S, Bo X and Yu G. clusterProfiler 4.0: A universal enrichment tool for interpreting omics data. Innovation (Camb). 2021;2:100141.

24. Bellini C, Bersi MR, Caulk AW, Ferruzzi J, Milewicz DM, Ramirez F, Rifkin DB, Tellides G, Yanagisawa H and Humphrey JD. Comparison of 10 murine models reveals a distinct biomechanical phenotype in thoracic aortic aneurysms. J R Soc Interface. 2017;14.

25. Bersi MR, Bellini C, Humphrey JD and Avril S. Local variations in material and structural properties characterize murine thoracic aortic aneurysm mechanics. Biomech Model Mechanobiol. 2019;18:203–218.

26. Humphrey JD, Schwartz MA, Tellides G and Milewicz DM. Role of mechanotransduction in vascular biology: focus on thoracic aortic aneurysms and dissections. Circ Res. 2015;116:1448–1461.

27. Shen YH, LeMaire SA, Webb NR, Cassis LA, Daugherty A and Lu HS. Aortic Aneurysms and Dissections Series: Part II: Dynamic Signaling Responses in Aortic Aneurysms and Dissections. Arterioscler Thromb Vasc Biol. 2020;40:e78–e86.

28. Yi X, Zhou Y, Chen Y, Feng X, Liu C, Jiang DS, Geng J, Li X, Jiang X and Fang ZM. The expression patterns and roles of lysyl oxidases in aortic dissection. Front Cardiovasc Med. 2021;8:692856.

29. Lee VS, Halabi CM, Hoffman EP, Carmichael N, Leshchiner I, Lian CG, Bierhals AJ, Vuzman D, Mecham RP, Frank NY and Stitziel NO. Loss of function mutation in LOX causes thoracic aortic aneurysm and dissection in humans. Proc Natl Acad Sci U S A. 2016;113:8759–8764.

30. Molnar J, Fong KS, He QP, Hayashi K, Kim Y, Fong SF, Fogelgren B, Szauter KM, Mink M and Csiszar K. Structural and functional diversity of lysyl oxidase and the LOX-like proteins. Biochim Biophys Acta. 2003;1647:220–224.

31. Maki JM, Rasanen J, Tikkanen H, Sormunen R, Makikallio K, Kivirikko KI and Soininen R. Inactivation of the lysyl oxidase gene Lox leads to aortic aneurysms, cardiovascular dysfunction, and perinatal death in mice. Circulation. 2002;106:2503–2509.

32. Hayashi K, Fong KS, Mercier F, Boyd CD, Csiszar K and Hayashi M. Comparative immunocytochemical localization of lysyl oxidase (LOX) and the lysyl oxidase-like (LOXL) proteins: changes in the expression of LOXL during development and growth of mouse tissues. J Mol Histol. 2004;35:845–855.

33. Guo DC, Regalado ES, Gong L, Duan X, Santos-Cortez RL, Arnaud P, Ren Z, Cai B, Hostetler EM, Moran R, Liang D, Estrera A, Safi HJ, Leal SM, Bamshad MJ, Shendure J, Nickerson DA, Jondeau G, Boileau C and Milewicz DM. LOX mutations predispose to thoracic aortic aneurysms and dissections. Circ Res. 2016;118:928–934.

34. Li H, Guo J, Jia Y, Kong W and Li W. LOXL4 abrogation does not exaggerate angiotensin II-induced thoracic or abdominal aortic aneurysm in mice. Genes (Basel). 2021;12.

35. Rodriguez HM, Vaysberg M, Mikels A, McCauley S, Velayo AC, Garcia C and Smith V. Modulation of lysyl oxidase-like 2 enzymatic activity by an allosteric antibody inhibitor. J Biol Chem. 2010;285:20964–20974.

36. Hajdú I, Kardos J, Major B, Fabó G, Lőrincz Z, Cseh S and Dormán G. Inhibition of the LOX enzyme family members with old and new ligands. Selectivity analysis revisited. Bioorg Med Chem Lett. 2018;28:3113–3118.

37. Kothapalli D, Liu SL, Bae YH, Monslow J, Xu T, Hawthorne EA, Byfield FJ, Castagnino P, Rao S, Rader DJ, Pure E, Phillips MC, Lund-Katz S, Janmey PA and Assoian RK. Cardiovascular protection by ApoE and ApoE-HDL linked to suppression of ECM gene expression and arterial stiffening. Cell Rep. 2012;2:1259–1271.

38. Anzai A, Shimoda M, Endo J, Kohno T, Katsumata Y, Matsuhashi T, Yamamoto T, Ito K, Yan X, Shirakawa K, Shimizu-Hirota R, Yamada Y, Ueha S, Shinmura K, Okada Y, Fukuda K and Sano M. Adventitial CXCL1/G-CSF expression in response to acute aortic dissection triggers local neutrophil recruitment and activation leading to aortic rupture. Circ Res. 2015;116:612–623.

39. Chen JY, Tsai PJ, Tai HC, Tsai RL, Chang YT, Wang MC, Chiou YW, Yeh ML, Tang MJ, Lam CF, Shiesh SC, Li YH, Tsai WC, Chou CH, Lin LJ, Wu HL and Tsai YS. Increased aortic stiffness and attenuated lysyl oxidase activity in obesity. Arterioscler Thromb Vasc Biol. 2013;33:839–846.

40. Kawai T, Takayanagi T, Forrester SJ, Preston KJ, Obama T, Tsuji T, Kobayashi T, Boyer MJ, Cooper HA, Kwok HF, Hashimoto T, Scalia R, Rizzo V and Eguchi S. Vascular ADAM17 (a Disintegrin and Metalloproteinase Domain 17) is required for angiotensin II/beta-aminopropionitrile-induced abdominal aortic aneurysm. Hypertension. 2017;70:959–963.

41. Sawada H, Ohno-Urabe S, Ye D, Franklin MK, Moorleghen JJ, Howatt DA, Mullick AE, Daugherty A and Lu HS. Inhibition of the renin-angiotensin system fails to suppress beta-aminopropionitrile-induced thoracic aortopathy in mice. Arterioscler Thromb Vasc Biol. 2022;42:1254–1261.

42. Wortmann M, Arshad M, Peters AS, Hakimi M, Böckler D and Dihlmann S. The C57Bl/6J mouse strain is more susceptible to angiotensin II-induced aortic aneurysm formation than C57Bl/6N. Atherosclerosis. 2021;318:8–13.

43. Henriques TA, Huang J, D’Souza SS, Daugherty A and Cassis LA. Orchidectomy, but not ovariectomy, regulates angiotensin II-induced vascular diseases in apolipoprotein E-deficient mice. Endocrinology. 2004;145:3866–3872.

44. Fashandi AZ, Spinosa M, Salmon M, Su G, Montgomery W, Mast A, Lu G, Hawkins RB, Cullen JM, Sharma AK, Ailawadi G and Upchurch GR, Jr. Female mice exhibit abdominal aortic aneurysm protection in an established pupture model. J Surg Res. 2020;247:387–396.

45. Chen JZ, Sawada H, Ye D, Katsumata Y, Kukida M, Ohno-Urabe S, Moorleghen JJ, Franklin MK, Howatt DA, Sheppard MB, Mullick AE, Lu HS and Daugherty A. Deletion of AT1a (Angiotensin II Type 1a) receptor or inhibition of angiotensinogen synthesis attenuates thoracic aortopathies in fibrillin1(C1041G/+) mice. Arterioscler Thromb Vasc Biol. 2021;41:2538–2550.

46. Robinet P, Milewicz DM, Cassis LA, Leeper NJ, Lu HS and Smith JD. Consideration of sex differences in design and reporting of experimental arterial pathology studies-statement from ATVB council. Arterioscler Thromb Vasc Biol. 2018;38:292–303.

47. Berman AG, Romary DJ, Kerr KE, Gorazd NE, Wigand MM, Patnaik SS, Finol EA, Cox AD and Goergen CJ. Experimental aortic aneurysm severity and growth depend on topical elastase concentration and lysyl oxidase inhibition. Sci Rep. 2022;12:99.

48. Fyfe FW, Gillman T and Oneson IB. A combined quantitative chemical, light, and electron microscope study of aortic development in normal and nitrile-treated mice. Possible implications for elucidating nitrile-induced aortic lesions and regarding the genesis of spontaneous arterial lesions. Ann N Y Acad Sci. 1968;149:591–627.

49. Qi X, Wang F, Chun C, Saldarriaga L, Jiang Z, Pruitt EY, Arnaoutakis GJ, Upchurch GR, Jr. and Jiang Z. A validated mouse model capable of recapitulating the protective effects of female sex hormones on ascending aortic aneurysms and dissections (AADs). Physiol Rep. 2020;8:e14631.

50. Nishida N, Aoki H, Ohno-Urabe S, Nishihara M, Furusho A, Hirakata S, Hayashi M, Ito S, Yamada H, Hirata Y, Yasukawa H, Imaizumi T, Tanaka H and Fukumoto Y. High salt intake worsens aortic dissection in mice: involvement of IL (interleukin)-17A-dependent ECM (extracellular matrix) metabolism. Arterioscler Thromb Vasc Biol. 2020;40:189–205.

51. Groth KA, Stochholm K, Hove H, Kyhl K, Gregersen PA, Vejlstrup N, Ostergaard JR, Gravholt CH and Andersen NH. Aortic events in a nationwide Marfan syndrome cohort. Clin Res Cardiol. 2017;106:105–112.

52. Trolle C, Mortensen KH, Hjerrild BE, Cleemann L and Gravholt CH. Clinical care of adult Turner syndrome--new aspects. Pediatr Endocrinol Rev. 2012;9 Suppl 2:739–49.

53. Eagleton MJ. Arterial complications of vascular Ehlers-Danlos syndrome. J Vasc Surg. 2016;64:1869–1880.

54. Hofmann CS, Wang X, Sullivan CP, Toselli P, Stone PJ, McLean SE, Mecham RP, Schreiber BM and Sonenshein GE. B-Myb represses elastin gene expression in aortic smooth muscle cells. J Biol Chem. 2005;280:7694–7701.

55. Murtada SI, Kawamura Y, Li G, Schwartz MA, Tellides G and Humphrey JD. Developmental origins of mechanical homeostasis in the aorta. Dev Dyn. 2021;250:629–639.

56. Owens APr, Subramanian V, Moorleghen JJ, Guo Z, McNamara CA, Cassis LA and Daugherty A. Angiotensin II induces a region-specific hyperplasia of the ascending aorta through regulation of inhibitor of differentiation 3. Circ Res. 2010;106:611–619.

57. Majesky MW. Developmental basis of vascular smooth muscle diversity. Arterioscler Thromb Vasc Biol. 2007;27:1248–1258.

58. Sawada H, Rateri DL, Moorleghen JJ, Majesky MW and Daugherty A. Smooth muscle cells derived from second heart field and cardiac neural crest reside in spatially distinct domains in the media of the ascending aorta-Brief Report. Arterioscler Thromb Vasc Biol. 2017;37:1722–1726.

59. Sawada H, Katsumata Y, Higashi H, Zhang C, Li Y, Morgan S, Lee LH, Singh SA, Chen JZ, Franklin MK, Moorleghen JJ, Howatt DA, Rateri DL, Shen YH, LeMaire SA, Aikawa M, Majesky MW, Lu HS and Daugherty A. Second heart field-derived cells contribute to angiotensin II-mediated ascending aortopathies. Circulation. 2022;145:987–1001.

60. Majesky MW, Horita H, Ostriker A, Lu S, Regan JN, Bagchi A, Dong XR, Poczobutt J, Nemenoff RA and Weiser-Evans MC. Differentiated smooth muscle cells generate a subpopulation of resident vascular progenitor cells in the adventitia regulated by Klf4. Circ Res. 2017;120:296–311.

61. Weiss D, Latorre M, Rego BV, Cavinato C, Tanski BJ, Berman AG, Goergen CJ and Humphrey JD. Biomechanical consequences of compromised elastic fiber integrity and matrix cross-linking on abdominal aortic aneurysmal enlargement. Acta Biomater. 2021;134:422–434.

62. Ferruzzi J, Bersi MR and Humphrey JD. Biomechanical phenotyping of central arteries in health and disease: advantages of and methods for murine models. Ann Biomed Eng. 2013;41:1311–30.

63. Caulk AW, Humphrey JD and Murtada SI. Fundamental Roles of Axial Stretch in Isometric and Isobaric Evaluations of Vascular Contractility. J Biomech Eng. 2019;141:0310081–03100810.

