## Supplemental Materials for "β-aminopropionitrile Induces Distinct Pathologies in the Ascending and Descending Thoracic Aortic Regions of Mice"

**Running Head:** BAPN-induced Aortopathies

*These authors contributed equally to this project.

**Corresponding Authors:**
Hong S. Lu
Saha Cardiovascular Research Center
741 South Limestone, BBSRB Room 249
Lexington, KY 40536


Alan Daugherty
Saha Cardiovascular Research Center
741 South Limestone, BBSRB Room 243
Lexington, KY 40536


**Table S1: BAPN-induced aortic rupture in 5 mouse strains**

| Mouse Strain | C57BL/6J | C57BL/6N | B6/129SF1 | 129X1 |
|---|---|---|---|---|
| **C57BL/6N** | P=0.2 | | | |
| **B6/129SF1** | P=0.001 | P=0.2 | | |
| **129X1** | P<0.001 | P<0.001 | P<0.001 | |
| **FVB** | P<0.001 | P<0.001 | P<0.001 | P=0.2 |

BAPN (0.5% wt/vol) was administered in drinking water for 12 weeks. Necropsy was performed to confirm aortic rupture. Survival curve is shown in Figure 1B. Log-Rank analysis was used to determine P values.

**Table S2. BAPN-induced aortic pathologies in 5 mouse strains survived for 12 weeks of BAPN administration**

| Mouse Strain | C57BL/6J | C57BL/6N | B6/129SF1 | 129X1 | FVB |
|---|---|---|---|---|---|
| **Mice survived (N/total N)** | 1/20 | 4/20 | 4/19 | 18/21 | 19/19 |
| **Ascending pathology (N/Survived)** | 1/1 | 2/4 | 2/4 | 2/18 | 2/19 |
| **Descending thoracic pathology (N/Survived)** | 1/1 | 0/4 | 4/4 | 7/18 | 2/19 |

BAPN (0.5% wt/vol) was administered in drinking water for 12 weeks. Gross pathologies were confirmed by in situ images shown in Supplemental Figure S2. N: number

### MAJOR RESOURCES TABLES

#### Animals (in vivo studies) – All mice were purchased from The Jackson Laboratory

| Mouse Strain | Sex | Strain # | Persistent ID/URL |
|---|---|---|---|
| C57BL/6J | Male & Female | 000664 | https://www.jax.org/strain/000664 |
| C57BL/6N | Male | 005304 | https://www.jax.org/strain/005304 |
| B6/129SF1 | Male | 101043 | https://www.jax.org/strain/101043 |
| 129X1 | Male | 000691 | https://www.jax.org/strain/000691 |
| FVB | Male | 001800 | https://www.jax.org/strain/001800 |

#### Primary Antibodies for Immunostaining

| Antibody | Vendor | Cat # | Working Concentration |
|---|---|---|---|
| Rabbit anti-smooth muscle α-actin | abcam | ab5694 | 2 μg/mL |
| Rabbit anti-CD68 (E3O7V) | Cell Signaling Technology | 97778 | 0.1 μg/mL |
| Rabbit nonimmune IgG | ImmunoReagents | Rb-003-V | Same as the antibody of interest (0.1 - 8.6 μg/mL depending on primary antibodies) |
| Rat monoclonal to CD31 [RM0032-1D12] | abcam | ab56299 | 1 μg/mL |
| Anti-Collagen I antibody | abcam | ab21286 | 8.6 μg/mL |
| Rabbit anti-CD3 antibody [SP7] | abcam | ab16669 | 0.1 μg/mL |
| Rabbit anti-CD19 (Intracellular Domain) (D4V4B) | Cell Signaling Technology | 90176 | 1 μg/mL |

#### Secondary Antibodies for Immunostaining

| Antibody | Vendor | Cat # | Note |
|---|---|---|---|
| ImmPress Goat Anti-Rabbit IgG | Vector | MP-7451 | No concentration information available; Ready-to-use (per manufacturer) |
| ImmPress Goat Anti-Rat IgG | Vector | MP-7444 | |

#### Primary Antibodies for Western blot

| Antibody | Vendor | Cat # | Working Concentration |
|---|---|---|---|
| Rabbit anti-Phospho-SMAD2 (Ser465/467) (138D4) | Cell Signaling Technology | 3108 | 0.1 μg/mL |
| Rabbit anti-Smad2 (D43B4) XP® | Cell Signaling Technology | 5339 | 0.3 μg/mL |
| Rabbit anti-Phospho-p44/42 MAPK (Erk1/2) (Thr202/Tyr204) | Cell Signaling Technology | 9101 | 0.2 μg/mL |
| Rabbit anti-p44/42 MAPK (Erk1/2) | Cell Signaling Technology | 9102 | 0.02 μg/mL |
| Mouse anti-βactin | Sigma-Aldrich | A5441 | 0.3 μg/mL |

#### Secondary Antibodies for Western blot

| Antibody | Vendor | Cat # | Working Concentration |
|---|---|---|---|
| Goat anti-Rabbit IgG Antibody | Vector | PI-1000 | 0.3 μg/mL |
| Goat anti-Mouse IgG Antibody | Sigma-Aldrich | A2554 | 0.3 μg/mL |

### Animal Study Information Following the ARRIVE Essential 10

#### Figure 1A

| Groups | Sex | Age (weeks)* | Number (Start) | Number (Termination) |
|---|---|---|---|---|
| 0.1% BAPN | Male | 3 | 10 | 10 |
| 0.3% BAPN | Male | 3 | 10 | 4 |
| 0.5% BAPN | Male | 3 | 10 | 2 |

#### Figure 1B

| Groups | Sex | Age (weeks)* | Number (Start) | Number (Termination) |
|---|---|---|---|---|
| C57BL/6J | Male | 3 | 20 | 1 |
| C57BL/6N | Male | 3 | 20 | 4 |
| B6/129SF1 | Male | 3 | 19 | 4 |
| 129X1 | Male | 3 | 21 | 18 |
| FVB | Male | 3 | 19 | 19 |

#### Figure 1C

| Groups | Sex | Age (weeks)* | Number (Start) | Number (Termination) |
|---|---|---|---|---|
| 3-week-old | Male | 3 | 38 | 1 |
| 4-week-old | Male | 4 | 15 | 8 |
| 26-week-old | Male | 26 | 15 | 15 |

#### Figure 1D

| Groups | Age (weeks)* | Number (Start) | Number (Termination) |
|---|---|---|---|
| Male | 4 | 24 | 14 |
| Female | 4 | 23 | 12 |

#### Figure 2B

| Groups | Total Number | Ascending Aortic Rupture | Descending Aortic Rupture |
|---|---|---|---|
| Male | 78 | 10 | 68 |
| Female | 21 | 3 | 18 |

#### Figure 3B, E, and F

| Groups | Sex | Age (weeks)* | Number (Start) | Number (Termination) |
|---|---|---|---|---|
| Vehicle (Male) | Male | 4 | 10 | 10 |
| BAPN (Male) | Male | 4 | 24 | 14 |
| Vehicle (Female) | Female | 4 | 10 | 10 |
| BAPN (Female) | Female | 4 | 23 | 12 |

#### Figure 3D, G, and H

| Groups | Sex | Age (weeks)* | Number (Start) | Number (Termination) |
|---|---|---|---|---|
| Young (Male) | Male | 4 | 15 | 8 |
| Mature (Male) | Male | 26 | 15 | 15 |
| Young (Female) | Female | 4 | 15 | 7 |
| Mature (Female) | Female | 26 | 15 | 15 |

**Figure 6A-C**

| Groups | Sex | Age (weeks)* | Number (Start) | Number (Termination) |
|---|---|---|---|---|
| Vehicle | Male | 4 | 20 | 20 (pooled 4 aorta/sample, so N=4 for RNA sequencing) |
| BAPN | Male | 4 | 20 | 20 (pooled 4 aorta/sample, so N=4 for RNA sequencing) |

**Figure 6D**

| Groups | Sex | Age (weeks)* | Number (Start) | Number (Termination) |
|---|---|---|---|---|
| Vehicle | Male | 4 | 5 | 5 |
| BAPN | Male | 4 | 5 | 5 |

**Figure 6E**

| Groups | Sex | Age (weeks)* | Number (Start) | Number (Termination) |
|---|---|---|---|---|
| Vehicle | Male | 4 | 5 | 5 |
| BAPN | Male | 4 | 5 | 4 |

**Figure S1**

| Mouse Strain | Sex | Age (weeks)* | Number (Start) | Number (Termination) |
|---|---|---|---|---|
| C57BL/6J | Male | 3 | 20 | 1 |
| C57BL/6N | Male | 3 | 20 | 4 |
| B6/129SF1 | Male | 3 | 19 | 4 |
| 129X1 | Male | 3 | 21 | 18 |
| FVB | Male | 3 | 19 | 19 |

**Figure S3A**

| Groups | Sex | Age (weeks)* | Number (Start) | Number (Termination) |
|---|---|---|---|---|
| Vehicle (Male) | Male | 4 | 10 | 10 |
| BAPN (Male) | Male | 4 | 24 | 14 |
| Vehicle (Female) | Female | 4 | 10 | 10 |
| BAPN (Female) | Female | 4 | 23 | 12 |

**Figure S3B**

| Groups | Sex | Age (weeks)* | Number (Start) | Number (Termination) |
|---|---|---|---|---|
| Young (Male) | Male | 4 | 15 | 8 |
| Mature (Male) | Male | 26 | 15 | 15 |
| Young (Female) | Female | 4 | 15 | 7 |
| Mature (Female) | Female | 26 | 15 | 15 |

**Figure S7**

| Groups | Sex | Age (weeks)* | Number (Start) | Number (Termination) |
|---|---|---|---|---|
| Vehicle | Male | 4 | 4 | 4 |
| BAPN | Male | 4 | 8 | 8 |

*age (weeks): the age when BAPN administration was started.

### ARRIVE Essential 10 Checklist

| Item | Application |
|---|---|
| Ethics | Approved by the University of Kentucky IACUC (2018-2967). |
| Sex | Described in each figure. |
| Inclusion criteria | Based on sex, age, body weight, and overt health appearance in each experiment. |
| Exclusion criteria | Based on sex, age, body weight, or medical cases reported by a veterinarian. |
| Sample size | Described in each figure legend. |
| Sample size calculation | None |
| Primary endpoint | Aortopathies |
| Randomization | Study mice were numbered and grouped randomly |
| Blinding | Quantification of aortic width (in situ) and luminal diameters (ultrasound) was measured by an investigator blinded to the study group information. Some data were measured by two investigators independently to validate the consistency of measurements. |
| Statistical analysis | SigmaPlot 15.0 (SYSTAT Software Inc., CA) or R Statistical Software. |
| Statistical method | Described in each figure legend |
| Data availability | All numerical data used for figures are available in the Supplemental Excel File. |

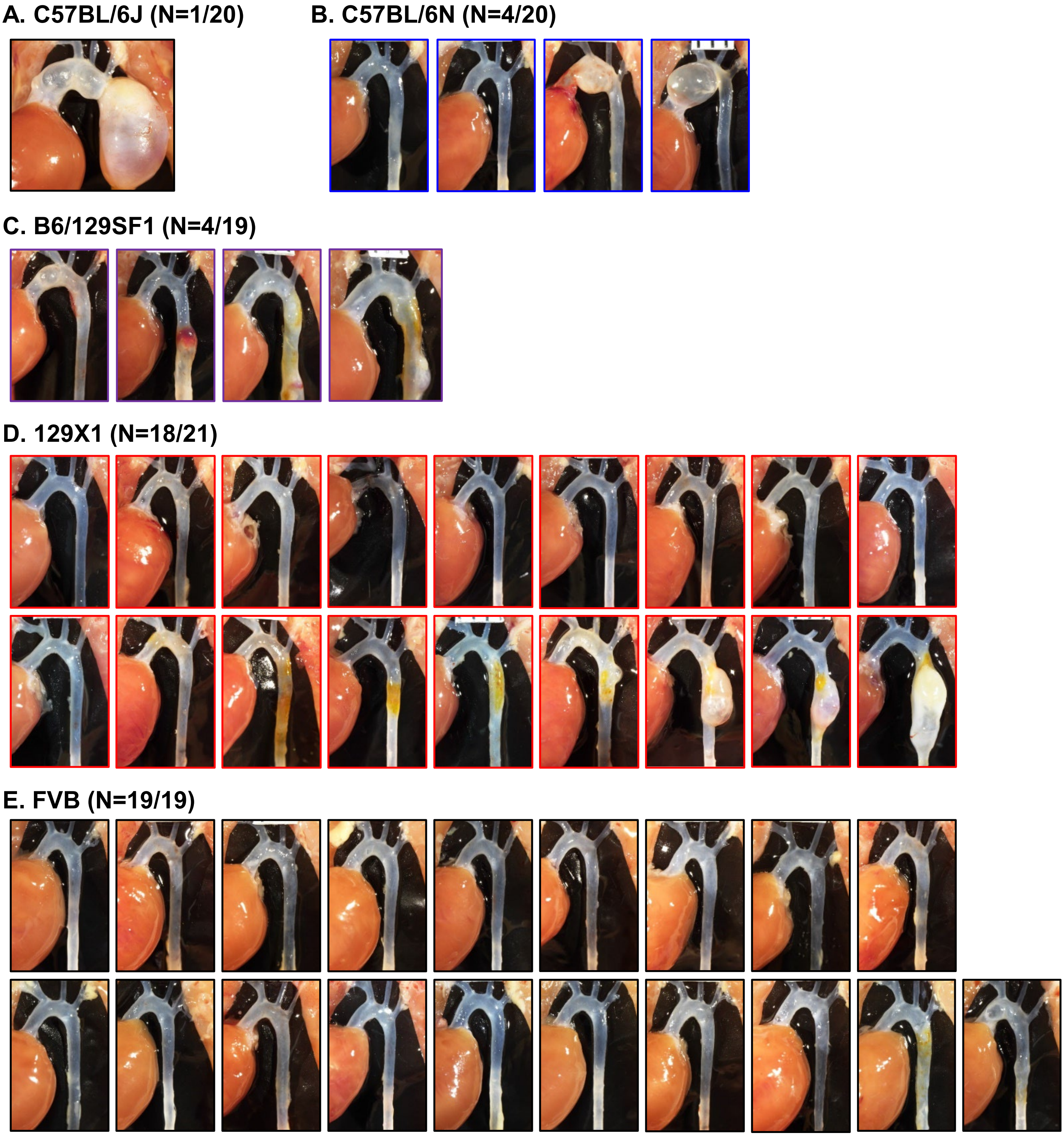


**Figure S1. BAPN-induced thoracic aortopathies in 5 mouse strains.** BAPN (0.5% wt/vol) was administered in drinking water for 12 weeks in 3-week-old male mice. In situ images of thoracic aortas for all survived mice with C57BL/6J **(A)**, C57BL/6N **(B)**, B6/129SF1 **(C)**, 129X1 **(D)**, and FVB **(E)** mouse strains after 12 weeks of BAPN administration.

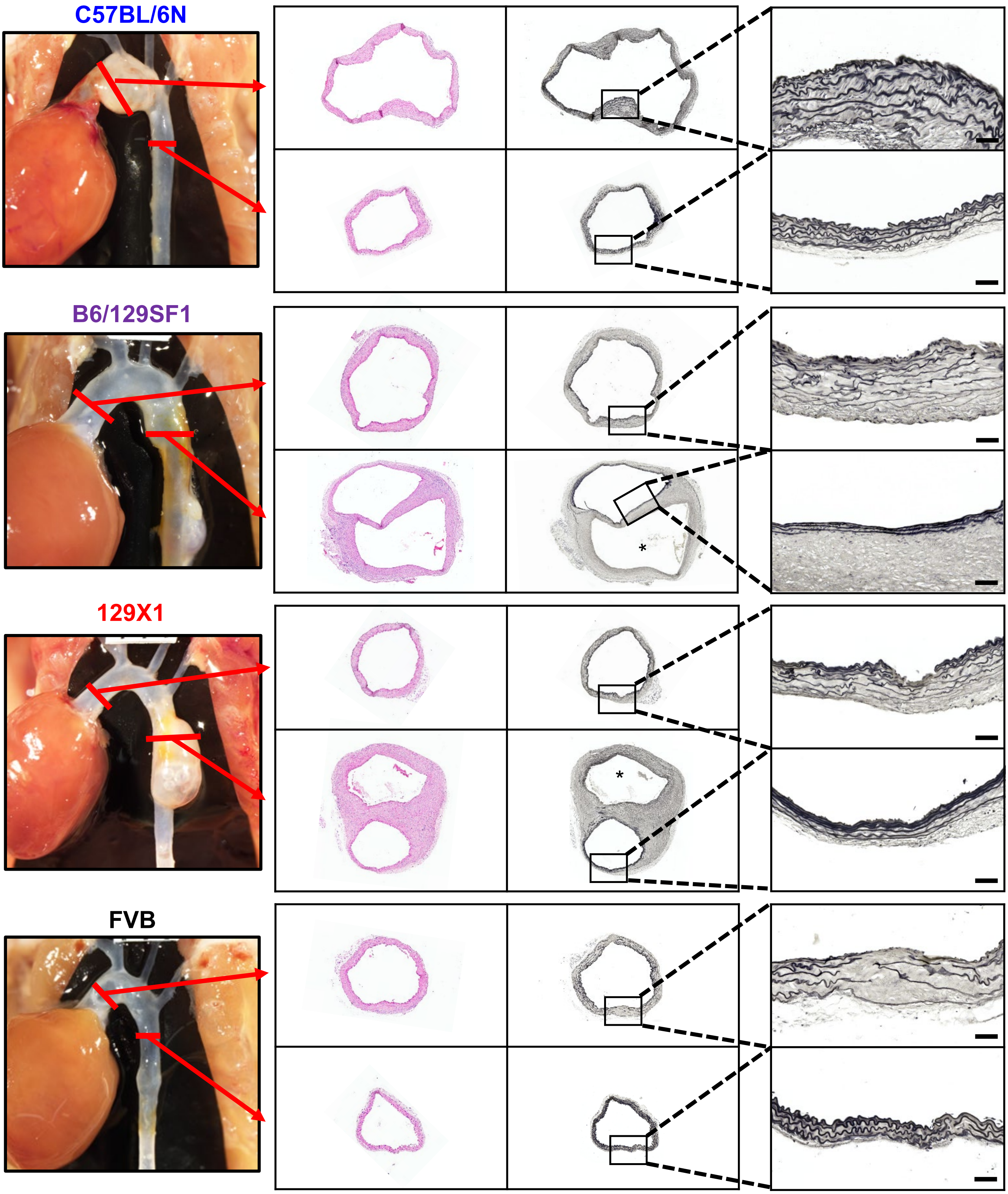


**Figure S2. BAPN-induced aortopathy in the ascending and descending thoracic regions had similar histological features across different mouse strains.** Representative images of hematoxylin-eosin (H&E) and Verhoeff's iron hematoxylin staining in ascending and descending thoracic aortic regions of mice administered BAPN for 12 weeks. The approximate locations of the tissue sections are indicated by red lines. * indicates the false lumen. Scale bar = 50 µm.

**A. 4-week-old**

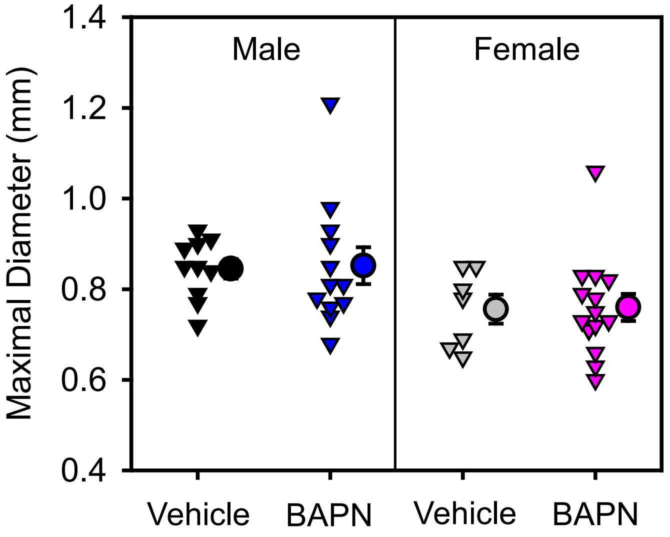


**B. Young (4-week-old) versus Mature (26-week-old)**

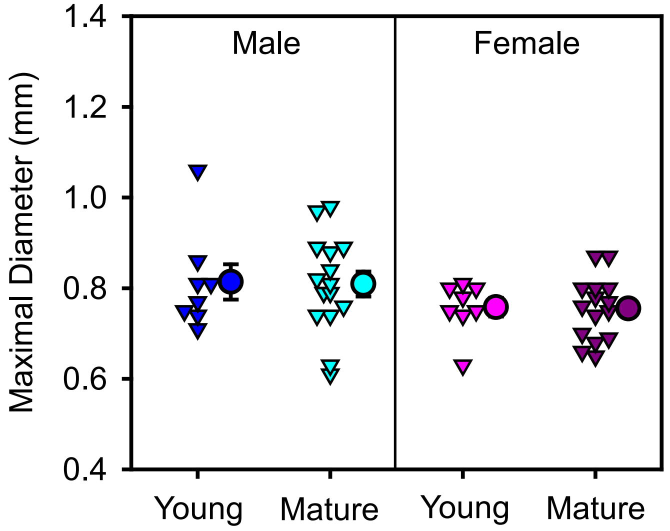


**Figure S3. BAPN did not induce dilatations of abdominal aortas in C57BL/6J mice.** Ex vivo measurements of maximal diameters of suprarenal aortas in 4-week-old C57BL/6J mice administered vehicle versus BAPN **(A)** and in 4- versus 26-week-old C57BL/6J mice administered BAPN **(B)** for 12 weeks.

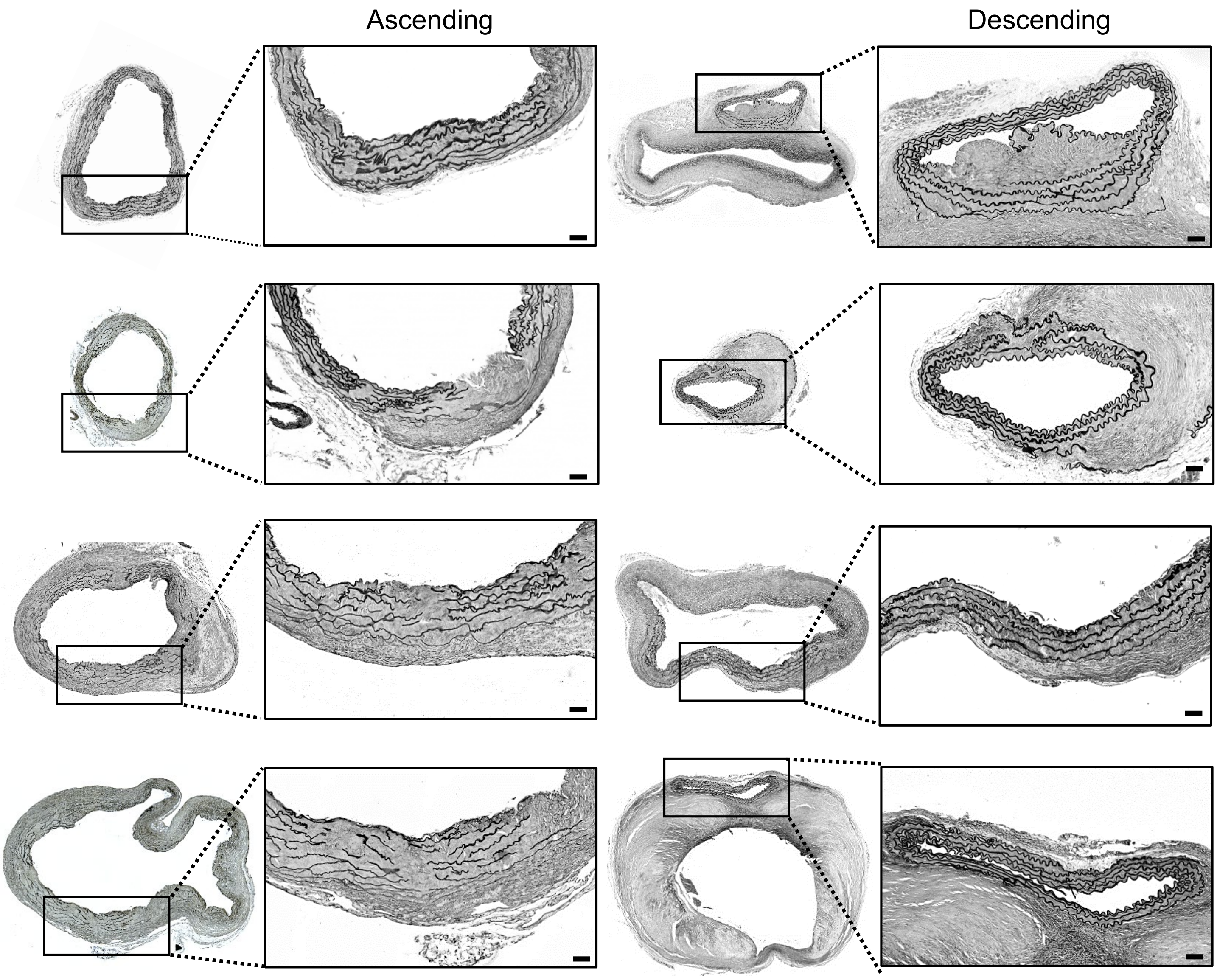


**Figure S4. BAPN induced extensive fragmentation of elastic fibers throughout the ascending aortic media, but not the descending thoracic aorta.** Representative images of Verhoeff staining in ascending and descending aortas from mice administered BAPN for 12 weeks. Scale bar = 50μm.

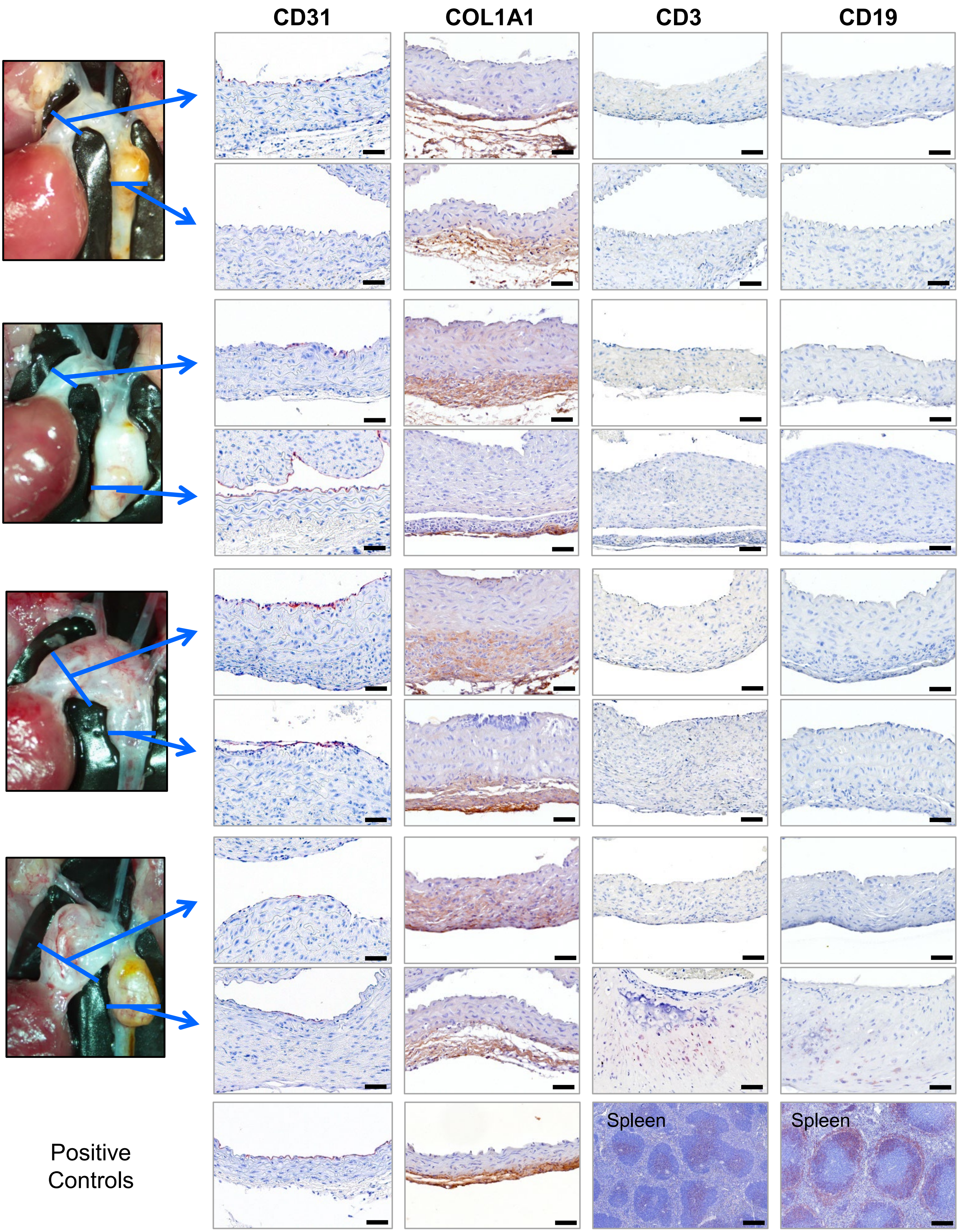


**Figure S5. Chronic administration of BAPN exhibited minimal CD3 or CD19-positive cell accumulation in the ascending and descending thoracic aortic regions.** Representative images of CD31, COL1A1, CD3, and CD19 immunostaining in ascending and descending aortas of mice with BAPN administration for 12 weeks. The approximate location of the tissue section from the ascending and descending region is indicated by the blue line. Scale bar = 50 μm for all aortas and = 200 μm for spleen.

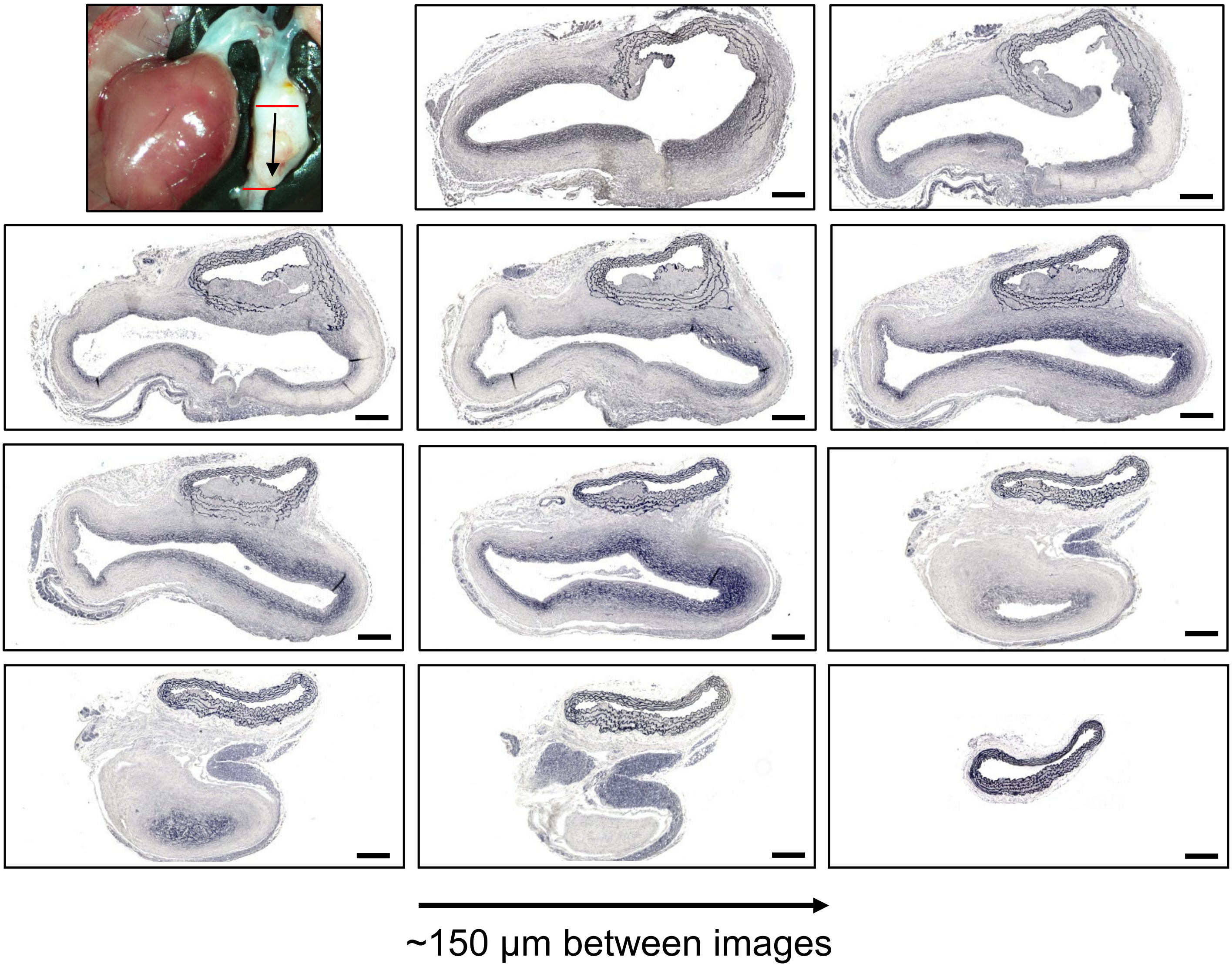


**Figure S6. Chronic BAPN administration led to complex pathological changes in the descending thoracic aorta of C57BL/6J mice.** BAPN was administered in drinking water for 12 weeks in a 4-week-old male C57BL/6J mouse. Histological images of Verhoff's iron hematoxylin staining from the descending thoracic aorta. Sections are presented from the proximal to distal region. Scale bar = 200 µm.

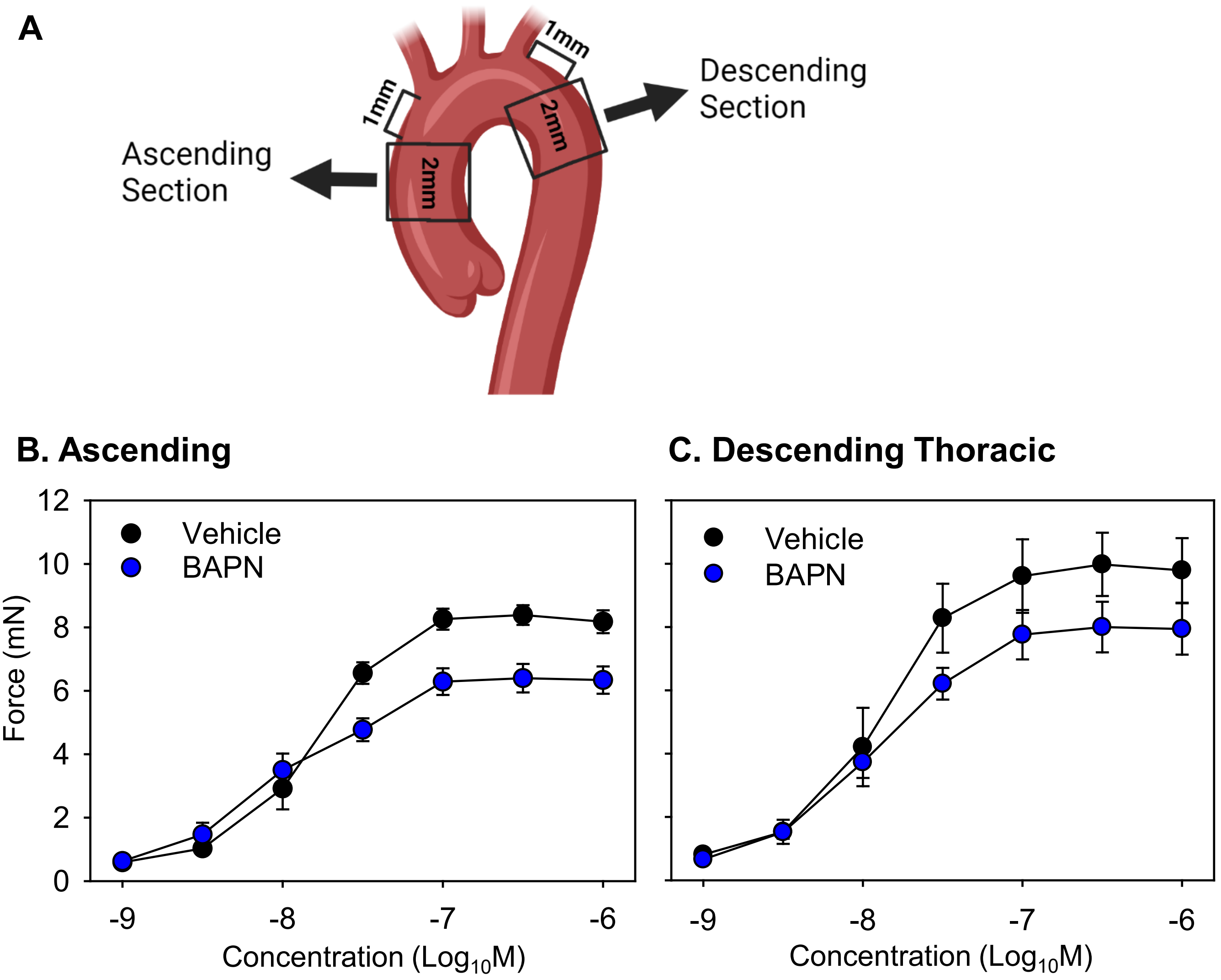


**Figure S7. BAPN decreased contractility in both the ascending and descending thoracic aortic regions.** Ascending and proximal descending thoracic aortas were harvested from 4-week-old male C57BL/6J mice administered either vehicle or BAPN for 7 days. **(A)** The cartoon shows the aortic regions where tissue was acquired for contractility analysis. The cartoon was created with BioRender.com. The contractile force in response to 5-HT (5-hydroxytryptamine) was measured in aortic rings harvested from **(B)** ascending and **(C)** descending thoracic regions. N=4 for vehicle and N=8 for BAPN group. Data were analyzed using Sigmoid function of R. $P<0.001$ and $P=0.04$ for comparing the inflection point values (distance from origin) between vehicle and BAPN groups in the ascending and descending thoracic aortic regions, respectively.
